# EiDA: A lossless approach for the dynamic analysis of connectivity patterns in signals; application to resting state fMRI of a model of ageing

**DOI:** 10.1101/2023.02.27.529688

**Authors:** Giuseppe de Alteriis, Eilidh MacNicol, Fran Hancock, Alessandro Ciaramella, Diana Cash, Paul Expert, Federico E. Turkheimer

**Affiliations:** Department of Neuroimaging, Institute of Psychiatry, Psychology and Neuroscience, King’s College London, London, UK; Global Business School for Health, University College London, London WC1E 6BT, UK; London Interdisciplinary Doctoral Programme, UCL Division of Biosciences, University College London, London WC1E 6BT, UK; Scuola Superiore Sant’Anna, Pisa, 56127, Italy

**Keywords:** Ageing, fMRI, Dynamic Functional Connectivity, Brain Network Dynamics, LEiDA, phase-locking

## Abstract

Dynamic Functional Connectivity (dFC) is the study of the dynamical patterns emerging from brain function. We introduce EiDA (Eigenvector Dynamic Analysis), a method that losslessly reduces the dimension of the instantaneous connectivity patterns of a time series to characterise dynamic Functional Connectivity (dFC). We apply EiDA to investigate the signatures of ageing on brain network dynamics in a longitudinal dataset of resting-state fMRI in ageing rats.

Previous dFC approaches have relied on the concept of the instantaneous phase of signals, computing the instantaneous phase-locking matrix (**iPL**) and its eigenvector decomposition. In this work, we fully characterise the eigenstructure of the **iPL** analytically, which provides a 1000 fold speed up in dFC computations.

The analytical characterization of the **iPL** matrix allows us to introduce two methods for its dynamic analysis. 1) Discrete EiDA identifies a discrete set of phase locking modes using k-means clustering on the decomposed **iPL** matrices. 2) Continuous EiDA provides a 2-dimensional “position” and “speed” embedding of the matrix; here, dFC is conceived as a continuous exploration of this 2-D space rather than assuming the existence of discrete brain states.

We apply EiDA to a cohort of 48 rats that underwent functional magnetic resonance imaging (fMRI) at four stages during the course of their lifetime. Using Continuous and Discrete EiDA we found that brain phase-locking patterns become less intense and less structured with ageing. Using information theory and metastability measures derived from the properties of the **iPL** matrix, we see that ageing reduces the available functional repertoire postulated to be responsible for flexible cognitive functions and overt behaviours, and reduces the area explored in the embedding space.

## 1 Introduction

The brain is recognized as a complex dynamic system ((1; 2; 3; 4)) whose activity is best characterized by patterns of interactions across its constituent parts, at multitude of scales, e.g. molecular, cellular, systemic ((5)). Consequently, dynamic functional connectivity (dFC) has become of major interest in the analysis of brain data in health or disease ((6; 7)), sourced from studies such as functional magnetic resonance imaging (fMRI) or electrophysiological recordings. dFC is an extension of functional connectivity (FC) ((8)), which aims to quantify the connectivity between signals in a functional sense, i.e., where the connections are not anatomically informed, but based on a measure of similarity of signal across the entire acquisition period ((9; 10)). Where FC is “static”, in the sense that it computes connectivity measures representative of the entire recording, for example using Pearson Correlation or Mutual Information, dFC includes the time evolution element in the analysis of connectivity patterns ((6; 7; 11; 12; 13)).

In pursuing this objective, any dFC approach faces at least two challenges. The first pertains to determining the appropriate size of the observation window for the desired connectivity matrix. To conduct a dFC analysis of *N* signals, where correlations matrices are computed based on a defined time window size, it becomes crucial to identify the ideal window size. Is there an optimal (heuristic or theoretical) method to ascertain this? (14). The second problem revolves around dimensionality. In the aforementioned example, we are confronted with the task of analyising the temporal evolution of an *N* × *N* matrix, which may not be symmetric and can present difficulties in terms of both analysis and lossless storage.

A powerful and commonly used solution for addressing the challenge of dimensionality is Leading Eigenvector Dynamic Analysis (LEiDA) ((15)). The Hilbert transform of the fMRI time-series is used to recover the analytical signal at each time point in the time-series ((16; 17)). Instantaneous amplitude and phase is extracted from the complex valued analytical signal, making it possible to measure pair wise phase differences ((18)). LEiDA computes an instantaneous Phase Locking Matrix (**iPL**) containing pairwise phase differences (e.g. phase locking). This matrix is then decomposed into its orthogonal components, the eigenvectors, where the first or leading eigenvector is selected and the remaining discarded, thus reducing the dimensionality from *N* × *N* to *N* × 1 dimensions. Clustering is then performed on the eigenvector time-series to interpret the temporal changes in the leading eigenvector. This enables the identification of distinct and reproducible spatiotemporal patterns, or “modes” (sometimes referred to as “brain states”) of phase-locking that the brain consistently exhibits throughout the recording period. The use of LEiDA has made important contributions to various functional brain paradigms, yielding insights across diverse areas of research, for example the study of sleep-wake transitions ((19)), action of psychedelic drugs ((20; 21)), neurodevelopmental paradigms ((22)), schizophrenia ((23)), and depression ((24; 25)).

Using the LEiDA approach, different synthetic dynamic indices of connectivity have been proposed such as the average duration of mode, or its fractional occurrence (the total occurrence of a cluster as a percentage of the full recording). In addition, within modes, it is possible to define measures capturing ideas drawn from complex systems theory such as metastability (a metric reflecting simultaneous tendencies for coupling and decoupling) ((26; 27; 28; 4)).

In this paper we introduce EiDA (Eigenvector Dynamic Analysis). We developed a closed form analytical decomposition of the **iPL** matrix and an approach that retains the complete information from the decomposition. Secondly, we investigated whether in certain circumstances it may be more appropriate to model dynamic connectivity as a smooth transition across FC configurations ((29)), instead of identifying, through clustering, discrete and separate brain modes.

In doing so, we identified the **iPL** matrix and its evolution as the most important and informative dFC object. Given the structure of the **iPL** matrix, it is always possible to decompose it analytically into two eigenvectors. This decomposition provides the theoretical background for **iPL**-based dynamic functional connectivity studies. Eigenvector decomposition allows the compression of the *N* × *N* **iPL** matrix into just 2*N* elements without loss and, furthermore, using the analytical form of the two eigenvectors drastically reduces their computation time (up to 1000x). We further demonstrated how the evolution of the two eigenvectors can be used to quantify the trajectories of the dynamic connectivity patterns. We use both a discrete state approach, using clustering of both the eigenvectors as in LEiDA, which we call Discrete EiDA, but also a continuous flow analysis using a 2-dimensional embedding, which we call Continuous EiDA. Together with EiDA, we propose two theoretically informed measures of phase locking based on the norm of the **iPL** matrix, namely the spectral radius and spectral metastability. All these measures are general enough to be applied to any dFC time-varying matrix, once the relevant number of eigenvectors is identified.

We applied EiDA to a longitudinal fMRI data-set acquired across the life-span of a cohort of rodents (four data-sets) (30; 31). This is a controlled experimental setting that challenges the methodology to recover the trajectories of expected loss of dynamical connectivity associated with ageing (29; 32; 33; 34; 35; 36; 37; 38; 39).

Previously, Betzel and co-workers demonstrated that, with ageing, functional connectivity within resting-state networks measured with fMRI decreases while functional connectivity between resting-state networks increases ((32)). In general terms, this highlights a loss of functional cohesion associated with the passing of time. These results were further expanded by the demonstration of reduced diversity/exploration of dynamic functional connectivity in fMRI data with age ((33)). Dynamic analysis of EEG data though has demonstrated somehow diverging results whereas the temporal complexity of phase synchronization was found to either increase, when measured with multiscale entropy,((35)) or decrease with age and this was associated with a slowing of the dynamic changes of the phase matrix ((29)).

## 2 Methods and Materials

A summary of our approach is outlined in Figure 1

**Figure 1.**
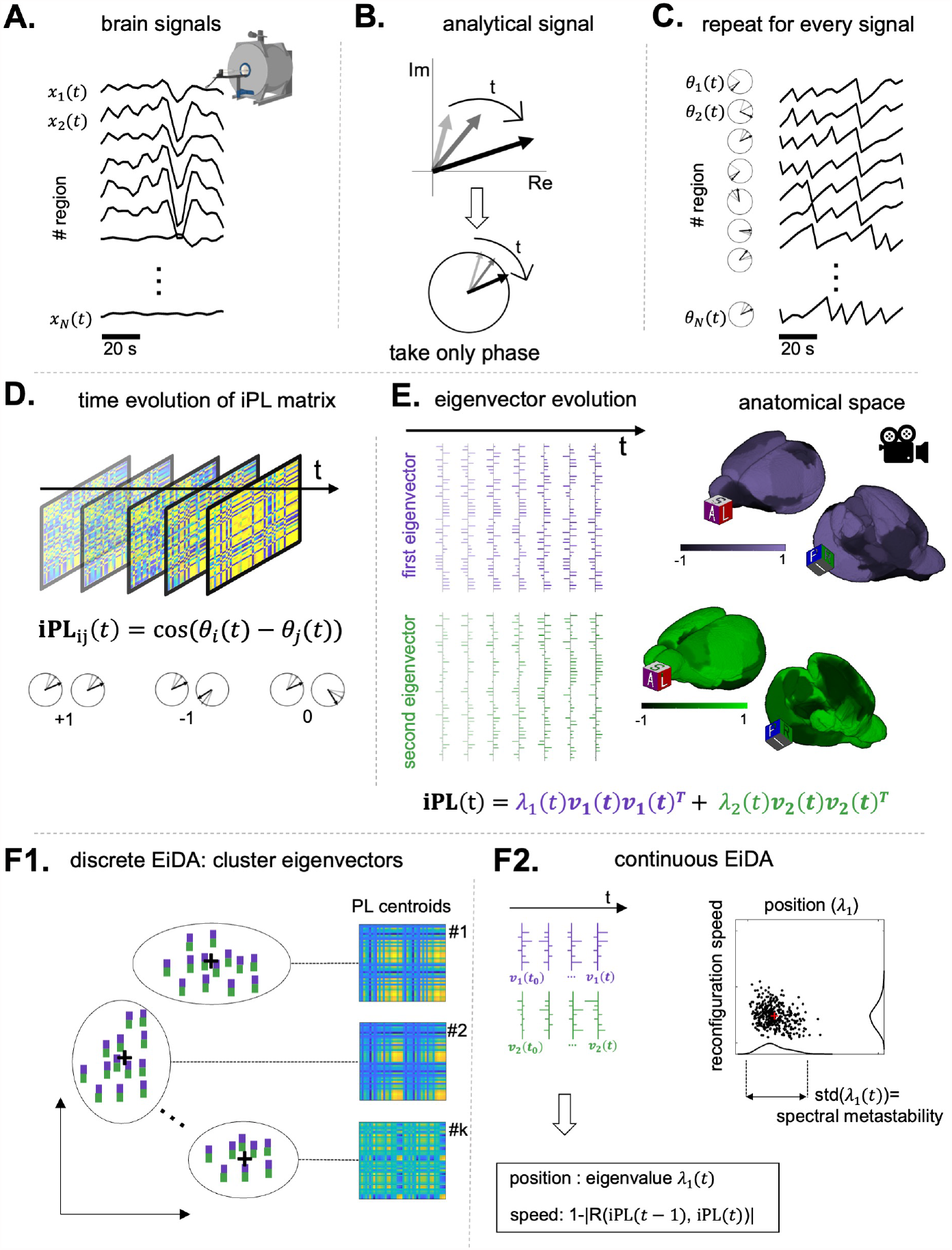
Summary of the proposed method. We started from a set of multi-dimensional time-series, the fMRI signals from 44 rat brain areas. **B**. For each signal, we computed its analytical representation via Hilbert Transform obtaining a complex number evolving in time, of which we considered only the phase. **C**. We repeated this procedure for every signal to obtain a multi-dimensional time-series of phases. **D**. At each time step, we computed the instantaneous Phase Locking Matrix (**iPL**), obtaining a time-series of matrices. The entry *i, j* of the matrix is the cosine of the difference of the phases *θ*_*i*_ and *θ*_*j*_. If the two phases are equal, the cosine of the difference is 1, -1 if they are opposite and 0 if they are in quadrature. **E**. The matrix being of rank 2 (see methods) we decomposed it into its two non trivial eigenvectors, therefore reducing the data to a timeseries of two vectors. Note that at each time *t*, the eigenvectors can be projected back in the anatomical space, as they have the same dimensionality of the timeseries. **F**. Once the timeseries of eigenvectors for each recording are obtained, we propose two alternative analysis strategies. *F1*. Discrete EiDA performs *k*-means clustering to identify *k* “modes”. See Section 2.8.1 for specifics about the algorithm. *F2*. Continuous EiDA, on the other hand, embeds the flow of phase-locking configurations in a 2D position-speed space, see Section 2.8.2. The first dimension is the first eigenvalue which is called “position”, because it is the norm of the **iPL** matrix. The second one is the speed at which the **iPL** matrix evolves, the reconfiguration speed. Alternatively, as in Figure 6, two position and speed plots can be computed using the two separate eigenvectors.

We defined a “recording” as the collection of *N* signals *x*_1_(*t*), *x*_2_(*t*), …, *x*_*N*_ (*t*), *t* = 1, …, *T*. We referred to a “group” as a set of recordings. In our case, as detailed in 2.11, the recordings are resting-state fMRI signals obtained during a two year study of brain ageing in rodents where the time-series in each recording were obtained from the parcellation of *N* = 44 anatomical brain regions ((30; 31)). The recordings were repeated for each animal four times during the life-span of the study. The study is described in section 2.11.

### 2.1 Static Analysis of recordings and Static Functional Connectivity

We used static connectivity measures, i.e. computed over the whole recording duration, ((40; 41)), as a benchmark and point of reference for the dynamic investigations of the recordings. The simplest measure we considered was the Matrix of Pearson Correlations (PC) of the paired time-series. For each recording, we defined a single matrix **P**, where **P**_*ij*_ = R(*x*_*i*_, *x*_*j*_), and R(.,.) is the Pearson correlation coefficient between the signals i and j. To show the effect of ageing in the static correlation patterns, an overall inter-subject connectivity matrix for each of the four groups of recordings was obtained by averaging the squared coefficient values in each group, thus obtaining four mean FC matrices. In the averaging process we calculated the squared values to take both positive and negative correlations into consideration. Additionally, we defined the FC Index as the sum of the squared values of the matrix ((42)), as a synthetic measure of overall connectivity for a single recording.

As already mentioned in the introduction, however, connectivity patterns may not be stationary over time. This means that the “static” connectivity matrix may not convey all the information about the dynamics of the connectivity patterns over time ((14; 43)). We then defined the **iPL** Matrix.

### 2.2 iPL Matrix

To perform a dynamic analysis and avoid the need to define time windows, one needs an instantaneous measure that can be used to compute the level of functional connectivity between each pair of brain regions. A common approach to this task is to obtain an analytical representation of a signal, which expresses a time series as a complex number, and thus an instantaneous amplitude *A*(*t*) and instantaneous phase *θ*(*t*). To compute the analytical signal, we used the Hilbert transform. The analytical form of a signal *x*(*t*) is equal to *x*(*t*) + *i*ℋ{*x*(*t*)}, where ℋ{*x*(*t*)} is the Hilbert transform of the signal. The Hilbert transform of a signal is defined as:

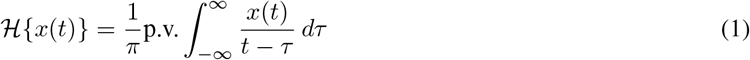

For more details about the Hilbert transform, please see ((44; 45)). To provide a visual illustration, (see Figure 1), we can conceive the analytical signal as a “clock” with the hand of the clock that changes length over time (with the amplitude *A*(*t*)) and rotates (changes with phase *θ*(*t*)). Therefore, at each time instant *t*, one could ask whether two signals are “phase locked”, i.e., they have the same instantaneous phase *θ*(*t*): in this case the two hands of the two clocks point in the same direction, (see Figure 1, C,D). This can be done for each pair of signals, and at each time point. We can thus define an “instantaneous” phase locking value **ipl**(**t**)_**1**,**2**_ between two signals *x*_1_(*t*) and *x*_2_(*t*) as the cosine of the difference of two phases **ipl**(**t**)_**1**,**2**_ = cos(*θ*_1_(*t*) − *θ*_2_(*t*)) ((44; 28; 15; 24; 25; 46; 26)). This value is equal to 1 if signals are perfectly in phase, -1 if their phase difference is *π*, and 0 if their phase difference is 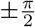, i.e. the signals are in quadrature.

It is therefore possible to define, given *N* signals *x*_1_(*t*), *x*_2_(*t*), …*x*_*N*_ (*t*), an instantaneous Phase Locking Matrix, **iPL** Matrix, i.e. a matrix where **iPL**_*ij*_(*t*) = cos(*θ*_*i*_(*t*) − *θ*_*j*_(*t*)) ((28)):

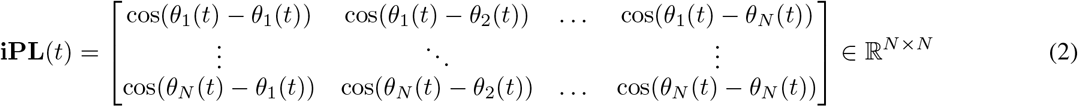

The analysis of connectivity patterns over time is transposed to the analysis of the evolution of the **iPL** matrix over time.

To compare static and dynamic connectivity matrix we computed the average **iPL** matrix for each recording, and then calculated the FC Index as in 2.1.

### 2.3 Analytical Computation of the Eigenvectors of iPL Matrix

Let us consider the *N* signals at a certain time *t* and their instantaneous phases, computed via Hilbert Transform, which form a vector 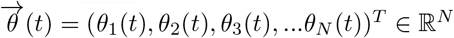. From this vector, we define, as in the previous section, an instantaneous Phase Locking (**iPL**) matrix, which is in principle different for each time *t*. Let us now, for simplicity, consider a single time *t*, and call the matrix **iPL** by abuse of notation, but remembering that the 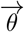 vector changes at every time *t* and so does the matrix. Hence, the next procedure can be repeated at each time *t*.

Given that the matrix **iPL**_*ij*_ = cos(*θ*_*i*_ − *θ*_*j*_), and given that cos(*x* − *y*) = cos(*x*) cos(*y*) + sin(*x*) sin(*y*), then **iPL**_*ij*_ = cos(*θ*_*i*_) cos(*θ*_*j*_) + sin(*θ*_*i*_) sin(*θ*_*j*_). This means that the *iPL* matrix can be decomposed in the sum of two matrices:

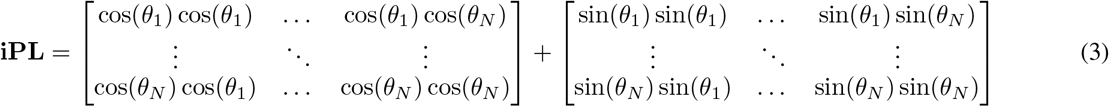

We define the two vectors, the “cosine” vector **c** = (cos(*θ*_1_), cos(*θ*_2_), …, cos(*θ*_*N*_))^*T*^ ∈ ℝ^*N*^, and the “sine” vector **s** = (sin(*θ*_1_), sin(*θ*_2_), …, sin(*θ*_*N*_))^*T*^ ∈ ℝ^*N*^, and rewrite the matrix as **iPL** = **cc**^*T*^ + **ss**^*T*^.

This matrix is symmetric and has ones on the diagonal, as **iPL**_*ii*_ = cos(*θ*_*i*_ − *θ*_*i*_) = cos(0) = 1. Its trace is thus *Tr*(**iPL**) = *N*. Importantly, this decomposition demonstrates that the matrix is a rank 2 matrix, which, being symmetric and positive semidefinite, will only have 2 non null (and greater than zero) eigenvalues *λ*_1_ and *λ*_2_ and their associated eigenvectors (as observed by (21)). This also implies that, as the sum of the eigenvalues of a matrix is equal to the trace of the matrix, *Tr*(**iPL**) = *N* = *λ*_1_ + *λ*_2_. Moreover, the two non-trivial eigenvectors will be a linear combination of **c** and **s**, so they will be of the form *A***c** + *B***s**. Thus, we only need to compute the two scalar values *A* and *B* to find the eigenvectors. Once we eigenvectors are found, we can impose the eigenvector equation to find the eigenvalues. See 5.1 and 5.2 for the full derivation.

Let us define the following quantities, which are the squared euclidean norm of the “cosine” and “sine” vectors and their scalar products:

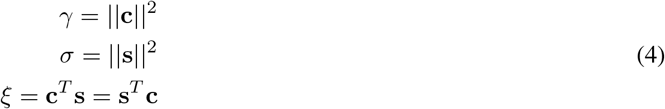

Based on these quantities, let us define:

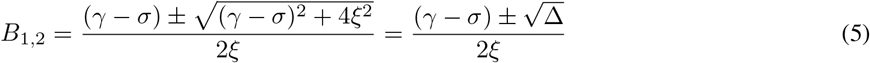

The two eigenvectors **v**_**1**_ and **v**_**2**_ are:

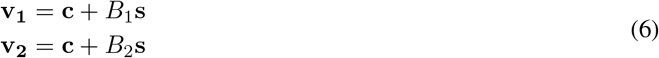

And their relative two non-null eigenvalues are:

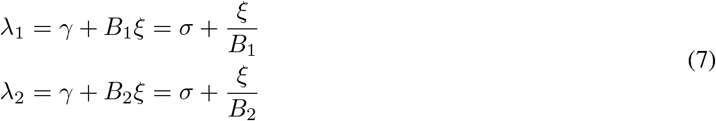

### 2.4 Two cases of interest

We now consider two limit cases, both of interest to understand the **iPL** matrix structure and the meaning of the eigenvalues.

The first case is *λ*_1_ = *N*, which implies *λ*_2_ = 0. The **iPL** matrix then has rank 1, as it only has 1 non null eigenvalue, see Figure 2, panel A. In this case, the **c** vector is parallel to the **s** vector. A rank 1 matrix with ones on the diagonal must have all elements equal to plus or minus one, i.e., this is the trivial case where all signals are in phase or antiphase: 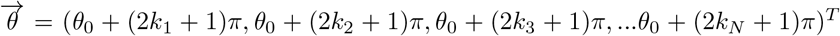, *k*_1…*N*_ ∈ ℤ. Then the matrix is maximally rank deficient and contains minimal information. Physiologically, this is the case of maximally phase locked signals, which occurs as the first eigenvalue *λ*_1_ tends to *N*.

**Figure 2.**
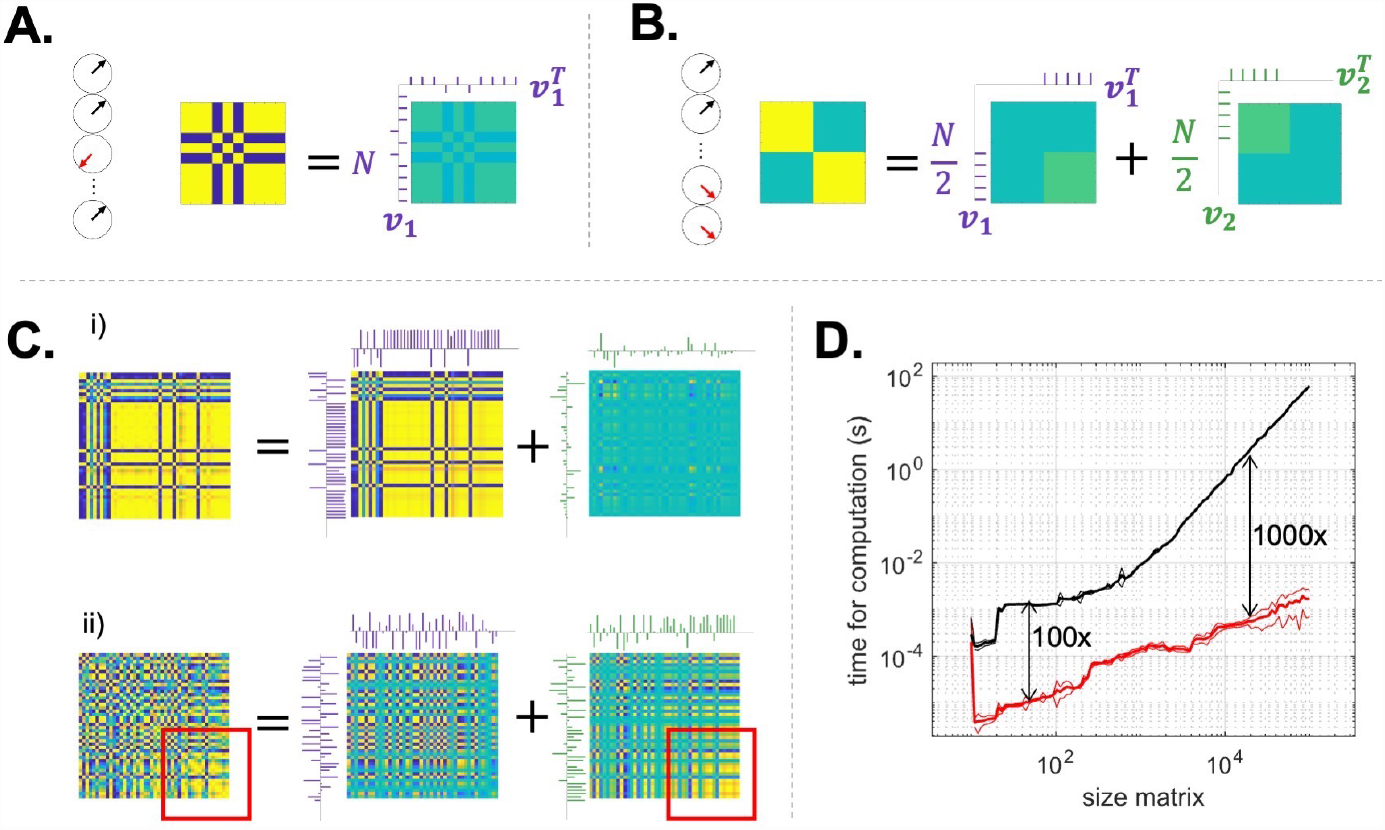
Illustration of the eigenvalue decomposition of limit cases matrices in **A** and **B** corresponding to *λ*_1_ = *N* and *λ*_1_ = *λ*_2_, and instructing exemplars from the rat data in **C**. Purple and green ticks indicate the first and second eigenvectors respectively. **A**. Rank 1 matrix: all signals are either in phase or in anti-phase, as indicated by the clocks. The matrix can be decomposed into a single eigenvector, with maximum eigenvalue *λ*_1_ = *N*. **B**. Both eigenvectors are equally important: the first N/2 signals are in quadrature with the second N/2. The two blocks of the matrix are reconstructed by summing 50% of the first eigenvector and 50% of the second. The eigenvalues are both equal to N/2. In this case, discarding one eigenvector would result in losing half of the information contained in the iPL matrix. **C**. Two interesting cases from our data. In the first one (i), the value of the first eigenvector is high, *λ*_1_ = 39 out of a maximum of 44, and contains most of the information. It is noticeable that in this case almost all signals are either in phase or in anti-phase, as can be seen from the large blocks. In the second case (ii), *λ*_1_ = 23, close almost N/2=22. In this case, the two eigenvectors contain almost the same amount of information. Throwing away the second one would lead to a large error in the reconstruction of the matrix: the information on the in phase hub highlighted by the red square is indeed contained in the second eigenvector. **D**. Average speed in seconds (mean, continuous line, plus or minus one standard deviation, dotted line) of our algorithm (red line) and the gold standard algorithm for iPL analysis (LEiDA), as a function of the size of the **iPL** matrix in a log log plot. Black arrows indicate that the black line is 100 or 1000 times the red one.

**Figure 3.**
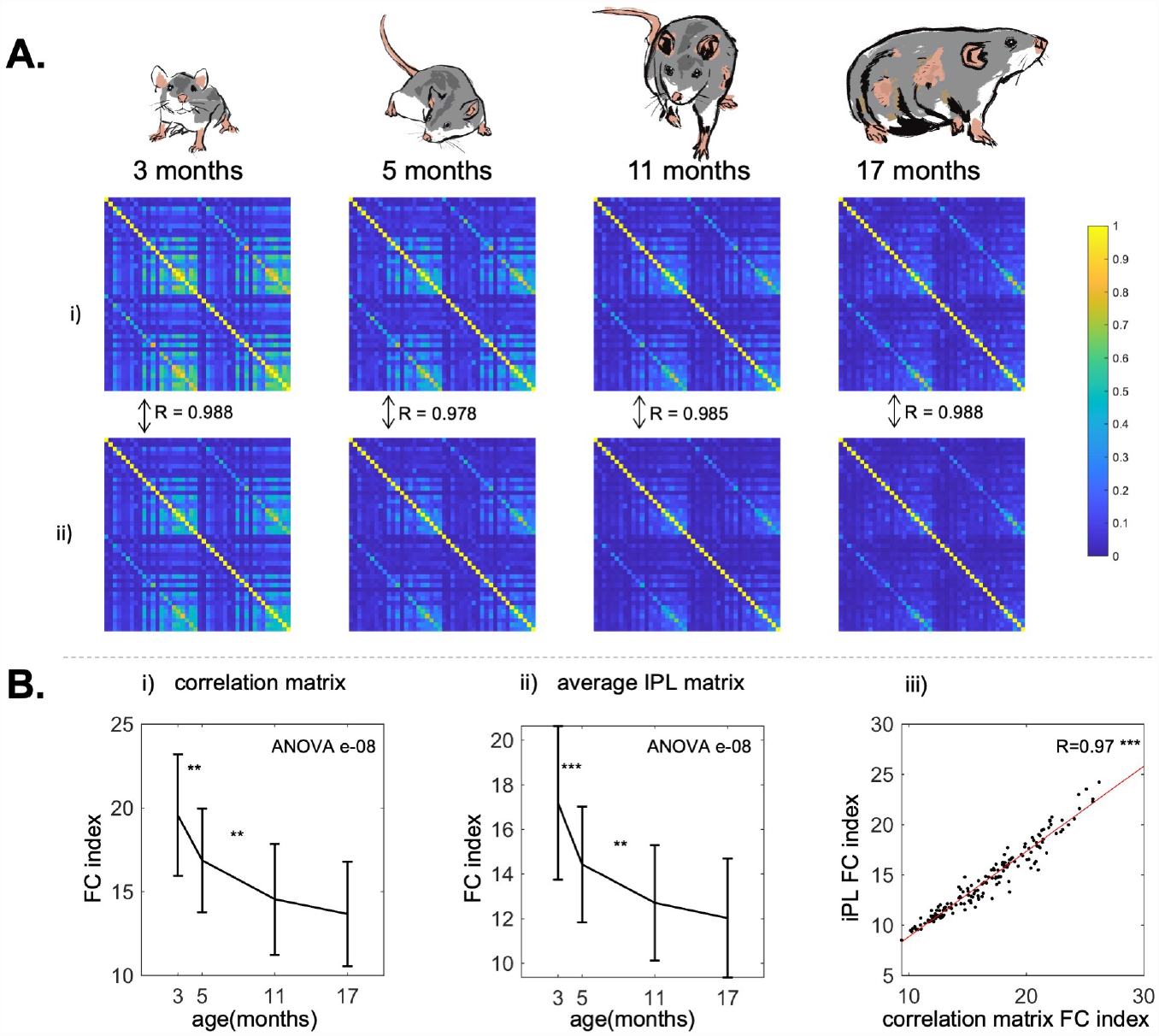
Results from the static analysis **A**. The group averaged Pearson connectivity matrix (i) and time averaged iPL matrix (ii) in the four age groups We indicate the element-wise correlations between the Pearson correlation matrices and the iPL matrices. Ageing is associated with an overall decrease of connected hubs (yellow hubs). **B**. The evolution with ageing (mean +-standard deviation) of the FC Index (see Section 2.1) for Pearson correlation matrices (i) and for the averaged iPL matrices (ii). Both the measures confirm a decrease which demonstrates the loss of connectivity strength observable in panel A. (iii): Correlation between the two measures with the best Ordinary Least Squares fit. The two measures are highly correlated, suggesting that the average iPL matrix conveys the same information as the Pearson correlation matrix.P-value of the nonparametric ANOVA test is reported inside the figures. * indicates p<0.05 ** p<0.01 and *** p<0.001 in the paired Wilcoxon tests between adjacent time points. Asterisks are reported only if p-values pass the Benjamini-Hochberg correction.

The second case is when *λ*_1_ = *λ*_2_ = *N/*2, see Figure 2, panel B. Given the constraints on the **iPL** matrix, this is possible if and only if **c** and **s** are orthogonal, their norms are equal and both equal to *N/*2, which is proven as follows. Given that the eigenvalues are *γ* + *B*_1_*ξ* and *γ* + *B*_2_*ξ*, and *γ* ≠ 0, by imposing *B*_1_ = *B*_2_ we have Δ = 0 and therefore *ξ* = 0 and *σ* = *γ*. This means that **c** and **s** are orthogonal and they have same norm, which is *γ* = *σ* = *N/*2 = *λ*_1_ = *λ*_2_. Note that if they are orthogonal it means that the two eigenvectors are **c** and **s** themselves, because **iPLc** = (**cc**^*T*^ + **ss**^*T*^)**c** = **cc**^*T*^ **c** = *γ***c** and similarly **iPLs** = *σ***s**.

Therefore, this is the case where the information contained in the **iPL** matrix is maximally irreducible to a single eigenvector, and therefore there is no “leading” eigenvector as the connectivity pattern is fully expressed by two orthogonal components which are both equally important, as the relative magnitude of the eigenvalues represents their contribution to the total information contained in the iPL matrix. An example of this configuration is a four blocks matrix where the two diagonal blocks are ones, all signals in phase, and the two non-diagonal blocks are zero, all signals in quadrature, see Figure 2, panel B.

Given these considerations on the **iPL** matrix, we define the following measures:

### 2.5 First eigenvalue = Spectral Radius

The **iPL** matrix is uniquely characterised by its first eigenvalue *λ*_1_, as *λ*_2_ = *N* − *λ*_1_ and all the other eigenvalues are null. *λ*_1_ is called the **spectral radius** of the **iPL** matrix, which corresponds to its 2-norm as the matrix is symmetric. Hence, it is a closed form norm of the instantaneous phase locking matrix ((42)). Indeed, as it tends to *N*, this norm indicates that all signals tend to the “trivial” state where they all are in phase or antiphase (See Section 2.4). At the same time, the matrix loses information and complexity by becoming maximally rank deficient and so maximally ordered. On the other hand, the more the norm approaches *N/*2, the less the matrix is irreducible to a single connectivity pattern and therefore one sees a less structured phase locking pattern.

### 2.6 Spectral Metastability

The spectral radius can be considered a global information metric about the instantaneous phase locking of the signals, expressed in closed form, fast to calculate, and conceptually similar to the Kuramoto Order Parameter ((28; 2; 26; 23; 47; 20; 48; 49)); note that the latter, however, is based on a specific model of the structure of the phase interactions, that is mean phase, while the first eigenvalue of the **iPL** matrix is general. Therefore, we can define a new general measure of metastability, the standard deviation over time of *λ*_1_, which is also equal to the standard deviation of *λ*_2_ as *λ*_2_ = *N* − *λ*_1_. We call this measure “spectral metastability” as the first eigenvalue is the spectral radius of the **iPL** matrix.

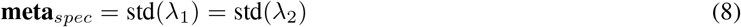

### 2.7 Irreducibility Index

The total information in the matrix is the sum of two orthogonal components, the eigenvectors, scaled by the two eigenvalues. If the first eigenvalue is lower than a percentage of *N* = *λ*_1_ + *λ*_2_, then this means that the reduction of the **iPL** matrix to the first eigenvector would keep less than this percentage of the total information. Firstly, we need to define a threshold representing the minimum amount of information one wants to be explained if we used only the first eigenvector whereas this threshold is expressed as a percentage of *N*. The Irreducibility Index is then the proportion of time during the recording in which the first eigenvalue of the **iPL** matrix is lower than the predefined threshold. If the experimenter requires to keep at least *x*% of the information at each time step, the Irreducibility Index at a level *x*% indicates the fraction of the recording in which reduction to the first eigenvector would fail in keeping this information. Thus a higher Irreducibility Index reflects a larger negative impact of the reliance on just the leading eigenvector.

### 2.8 Two new approaches to analyze eigenvector dynamics

The sections above demonstrated that the *N* × *N* **iPL** matrix can be fully and losslessly decomposed into two eigenvectors of size *N* and one eigenvalue. These form the basis for two approaches to analyze the dynamics of eigenvectors and eigenvalues over time. The first one is called “Discrete EiDA”, because it finds *k* discrete states by clustering the eigenvectors in line with the approach of the original LEiDA. The second is called “Continous EiDA” because it interprets the reconfiguration of eigenvectors as a continuous trajectory and quantifies its overall “position” and “speed” in a 2-D space. Based on dynamical considerations on the evolution of eigenvectors, as we will do in 3.2, the experimenter can choose to use the first or the second approach.

#### 2.8.1 Discrete EiDA

Discrete EiDA is a new k-means-based clustering algorithm that uses the information contained in the two eigenvectors to aggregate them in a discrete set of clusters that can be considered dFC-based brain states. Our proposed approach is mathematically equivalent to performing k-means with the full iPL matrices. However, it exploits the fact that the information of the matrix is fully contained in its two eigenvectors. Therefore, it operates on the eigenvectors instead of the full matrices allowing to compute k-means in an efficient way, and to store only the two eigenvectors instead of the full matrices (which may be computationally unfeasible). This is relevant as demonstrated in 3): in the data considered here, discarding the second eigenvector would neglect a significant amount of information, as indicated by the Irreducibility Index (defined in 2.4).

Each data-point entered into the algorithm is an **iPL** matrix, represented in a condensed form by its two non-trivial eigenvectors. The k-means clustering algorithm iterates between two steps: the first step assigns each data-point to one of k centroids by computing its distance from it. The second step updates the centroids by computing the mean of each cluster of data points.

This is the structure of the Discrete EiDA algorithm: the k centroids are saved as the centroid **iPL** matrices in the Upper Triangular form (computationally this is feasible, since k is a small number), while the data points are stored in a dataset *D* as stacked eigenvectors weighted with the square roots of eigenvalues. In the first step, all the distances are computed by rebuilding the matrix from the eigenvectors (**iPL** = *λ*_1_**v**_**1**_**v**_**1**_^*T*^ + *λ*_2_**v**_**2**_**v**_**2**_^*T*^) and computing the cosine distance between the upper triangular part of the rebuilt matrix and the centroid matrix. Then, in the second step, the centroid matrices are updated by averaging the upper triangular parts of the rebuilt matrices. Overall, we are clustering the full information contained in the matrices, without any loss, but we are using only the two non-trivial eigenvectors.

This is the pseudo-code for Discrete Eida:

##### Algorithm 1

Discrete EiDA K-means

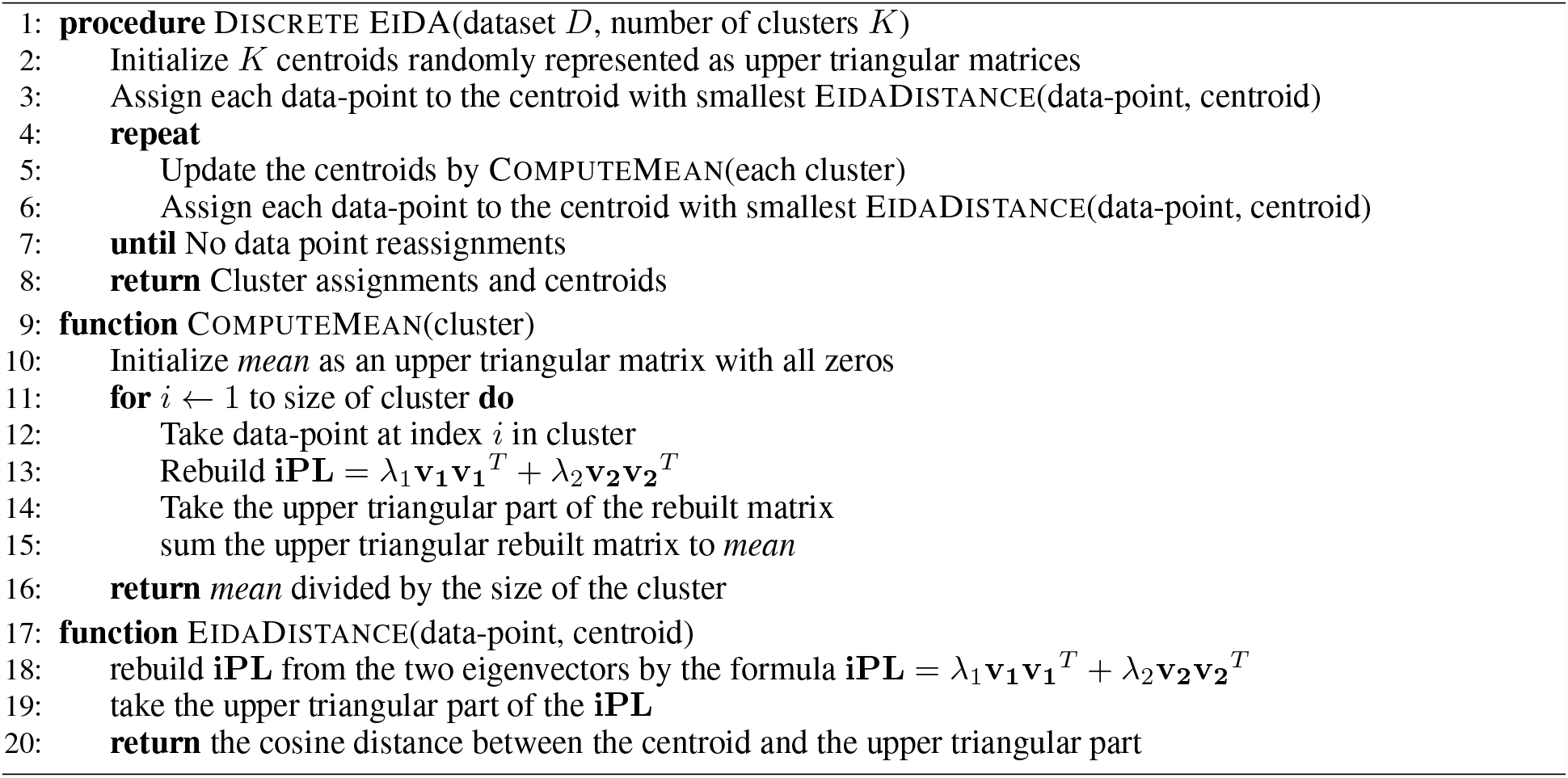

The final output of Discrete EiDA is *k* clusters with *k* centroids, which are identified as *k* phase-locking brain modes (they are the centroid **iPL** matrices). The clustering of the first eigenvectors was already proposed in previous approaches ((15; 26; 46; 20; 25; 24)). Here instead we propose to consider both eigenvectors because, as we have seen, the rank 2 **iPL** makes it possible to do clustering of the full information contained in the matrix in a lossless and efficient manner. Once the clusters are computed, it is then possible to visualize their representative states, the centroids, as averaged **iPL** matrices in the clusters. We highlight that since the centroids are the average of multiple rank-2 **iPL** matrices, their rank will no longer be 2. Indeed, they contain the full information on the average connectivity pattern in the clusters. Moreover, this new algorithm solves a known problem in the LEiDA literature, the flipping sign problem (23). Because eigenvectors have a sign ambiguity (an eigenvector is still an eigenvector if its sign is inverted) performing clustering in the leading eigenvector space poses a problem when two opposite sign eigenvectors representing the same state are clustered: in doing the mean of the clusters, they would cancel each other. Various approaches have been proposed to solve the flipping-sign problem ((21)) - however, we believe that with Discrete EiDA we solve the problem at the root by working in the native space of symmetric positive semidefinite **iPL** matrices, instead of the leading eigenvector space.

Moreover, once the clusters are computed, it is possible to use synthetic measures of the duration of modes (the same defined in LEiDA ((15; 46; 20; 25; 24))), in particular:

1. The Fractional Occurrence of a brain mode, i.e. the relative amount of time in the recording in which the **iPL** matrix belongs to a specific mode (=cluster).
2. The Dwell Time, i.e. the average duration of a mode (=cluster).

These two measures are not equivalent: a mode could appear very frequently in a recording, i.e. have a high fractional occurrence, but be on average for a very brief time: have a low dwell time).

Finally, within each cluster, it is possible to compute any dynamic measure of interest like spectral metastability (see Section 2.8), similarly to what done by (23), or to apply all the continuous measures introduced in Section 2.8.2.

Overall, we have described a specific way of performing clustering. The algorithm for Discrete Eida is available on GitHub at https://github.com/alteriis/EiDA.

However, we underline that a wide range of clustering methods exist beyond k-means clustering and might lead to different clusters, and they might be preferable depending on data, their distributions and the experimental questions.

Therefore, we propose that Discrete EiDA is a collection of clustering methods that use the eigenvector representation of the phase locking information to find a discrete set of brain modes.

It is important to underline that we do not expect the brain to present a clear and distinctive number of clusters (k), but to continuously explore its dynamic space. Hence we believe that a better conceptualization of the centroids is their use as masks to obtain time-varying **iPL** information metrics (like fractional occurrence, dwell time or metastability) rather than as real neurobiological states, particularly as they can be dependent on the specific clustering method used.

#### 2.8.2 Continuous EiDA

It is important to note that eigenvectors may not always be clustered into a a well-separated discrete set of centroids. This may manifest as a poor convergence of k-means and, equally, by the lack of spatial heterogeneity in the phase locking space/eigenspace exploration. In this case, the continuous exploration of Phase Locking configurations may be analyzed by plotting the dynamic walk of connectivity motifs in a 2D embedding ((50; 51)).

Therefore, we introduce an approach that examines the evolution of both eigenvectors over time as a continuous flow and call it Continuous EiDA.

To do so, we propose to follow the time evolution of **iPL** in a two dimensional “position-speed space”, using both eigenvectors in a kinematic speed-displacement (KSD) plot (52).

The two dimensions in this 2D embedding are: the “position” (=overall amount of connectivity at time *t*) *p*(*t*), and the “speed” of evolution of the **iPL** matrix *s*(*t*). The position is what the previous sections have demonstrated to be the best summary indicator of the state of the phase locking matrix: the spectral radius = first eigenvalue *λ*_1_. The speed is the “reconfiguration speed” (as already proposed by (29; 26)),at which the matrix evolves in its space, computed as the correlation distance between temporally adjacent matrices in their upper triangular form. We define reconfiguration speed *s*(*t*) as:

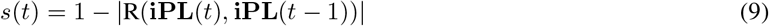

Where R is the Pearson Correlation Coefficient.

This is a 2-D embedding of the evolution of the **iPL** matrix. However, the individual evolution of each of the two eigenvectors may be also interesting to understand how the two orthogonal components of the **iPL** matrix change with time. It is possible to apply the same embedding for the two eigenvectors separately. In this case, the “position” will the the eigenvalue associated to the eigenvector (thus, *λ*_1_ for *v*_1_ and *λ*_2_ for *v*_2_) and the “speed” will be the same as in Equation 2.8.2 but computing the correlations of the eigenvectors instead of the full **iPL** matrix.

### 2.9 An alternative measure of phase synchronization

A commonly used measure of metastability can be obtained from one of the most popular measures of synchronisation: the Kuramoto Order Parameter

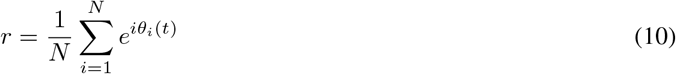

that can be computed for each time *θ*_1_(*t*), using the phases of the analytical signals((28; 2; 23; 26; 47; 20; 48; 49)). The modulus of *r* ∈ [0, 1] is a measure of synchronization: if all signals are in phase, this number approaches 1, while, if they are distributed uniformly, it will approach 0. It can be conceived as the mean of instantaneous phases. Metastability is defined as the standard deviation of the modulus of the Kuramoto order parameter over time ((28; 2; 23; 26; 47; 20; 48; 49)), and we refer to it as Kuramoto metastability to clearly differentiate it from spectral metastability introduced above.

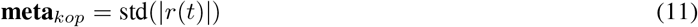

### 2.10 Informational Complexity

In order to have a proxy for the informational complexity of the evolution of the eigenvectors, we computed the number of bits required to compress it using the lossless data compression Lempel–Ziv–Welch (LZW) algorithm ((53; 54)). A higher complexity, reflected by the need to use more bits to store information, indicates more random and unpredictable patterns as there is more information to store, while a lower one indicates more coherent and structured patterns of evolution with less information to store. Similar approaches have been introduced to quantify perturbational complexity of EEG responses to Transcranial Magnetic Stimulation (TMS) ((55)).

### 2.11 Application of EiDA to rat fMRI data

We applied the methods described above to a longitudinal dataset of ageing rats ((31; 30), MacNicol et al, in preparation). A cohort of 48 Sprague Dawley rats (Charles River, UK) were monitored across their lifespan and scanned up to 4 times with a 9.4 T Bruker Biospec MR scanner, specifically at 3, 5, 11 and 17 months of age. These ages represent, respectively, late adolescence, young adulthood, middle age, and the beginning of senescence ((56)). Experiments were performed in accordance with the Home Office (Animals Scientific Procedures Act, UK, 1986) and approved by the King’s College London’s ethical committee. Resting-state functional data were recorded using a 2D multi-slice multi-echo echo planar imaging sequence with TR = 2750 ms, TEs = 11, 19, 27, and 35 ms, and a 70^*o*^ flip angle, producing an image with 40 slices. Slices were 0.5 mm thick with a 0.2 mm gap, which gave a 48 × 44 matrix, with an in-plane resolution of 0.5 × 0.5 mm. Rats were anesthetized with ca. 1.8% isoflurane for the duration of functional scans. This dose produced anatomically-plausible components from single-subject and group-level Independent Component Analysis (ICA) ((57; 58).

Motion-correction was estimated on the first echo time to its middle volume, and applied identically to each echo time volume. The corrected echo volumes were optimally combined, which maximises the signal-to-noise ratio at the expense of some loss of time resolution ((59)). Signals were simultaneously filtered with a 0.01-0.08 Hz band-pass filter and regressed for nuisance factors that included motion parameters and the CSF signal. Corrected and filtered fMRI volumes were warped to a study-specific template ((30))and parcellated into 44 anatomical regions of interest (ROI), 22 for each hemisphere, generated by combining delineations of predominantly gray matter structures from two popular rat atlases ((60; 61)). The BOLD signals were averaged within each ROI.

### 2.12 Statistical Analysis

We restricted our analyses to the 30 out of 48 rats that were scanned the maximum of four times. As measures were not normally distributed, we first tested the variation across age groups of all the parameters considered using a non-parametric one way ANOVA test (Kruskal-Wallis test). A post-hoc Wilcoxon rank sum test was then used to test the variability between each pair of consecutive age groups. The multiple comparison correction was performed by controlling the False Discovery Rate (FDR) at a rate *α* = 0.05, using the Benjamini-Hochberg procedure (62). The detailed p-values and ANOVA tables for all the statistics are reported in the Supplementary Material.

A Pearson Correlation Coefficient *R* was used to test collinearity between measures where the test for significance was obtained by calculating an empirical null *R* distribution by shuffling data.

## 3 Results

### 3.1 Static Functional Connectivity

A loss of total correlation and a diminution of the number of correlated areas over time is visible in the mean FC matrices for both PC and **iPL** connectivity (3 A).The FC Index (see Section 2.1) sharply decreases from 19.7 ± 3.9 at month 3 to 13.7 ± 3.1 at month 17 with an overall significance of e-08 (nonparametric ANOVA) in PC connectivity, with similar statistically significant decreases (ANOVA p=e-08) for iPL. The correlation between the FC Indexes of PC and iPL connectivity was both statistically significant (p<0.001) and high (R=0.97).

### 3.2 Dynamic Functional Connectivity measures: Discrete EiDA

We performed Discrete EiDA clustering with *k* (number of clusters) from 1 to 10. We plotted the sum of squared distances of each data point in a cluster with their centroids as a function of *k* to obtain the “elbow plot” (see Figure 4 C.) The point where this curve presents an elbow is often chosen as the optimal number of clusters because it indicates that adding another cluster (*k* = *k* + 1) would not significantly improve the clustering performance. We found k=3 as a satisfactory partitioning of the space. As specified in Section 2.8.1, we interpret the centroids of the spatiotemporal patterns as masks to analyze the dynamic reconfiguration of phase locking patterns and to compute dynamic measures like dwell time, fractional occurrence or metastability ((26)). As such we apply clustering for interpretation and quantification of brain network dynamics.

**Figure 4.**
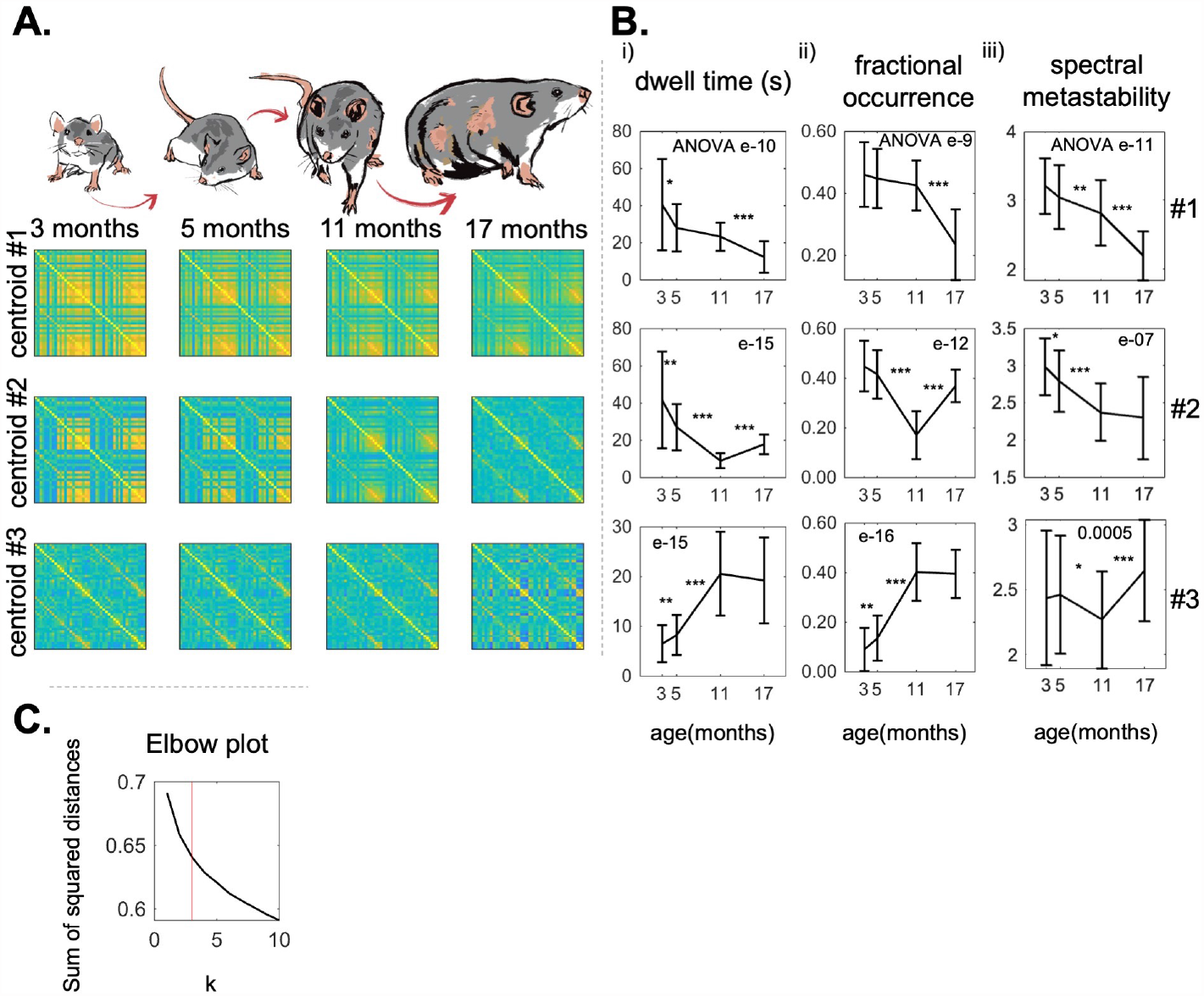
Results for the Discrete EiDA **A**: The three EiDA cluster centroids and their evolution with ageing. Each row represents a centroid, each column a different age group **B**: the evolution with ageing of (i) dwell time and (ii) fractional occurrence of clusters. The first two clusters, that are associated with connected hubs and reflect the connectivity information contained in the Pearson correlation matrices, become less frequent in the recordings as rats age. On the other hand, the third cluster, which is associated with a less structured and weaker connectivity pattern, increases in both dwell time and fractional occurrence with ageing. (iii) Spectral metastability in the three clusters. This measure decreases with ageing in the first two clusters, while there is not a clear pattern in the third. **C**. The elbow plot: the x axis is *k*, the y-axis is the the sum of squared distances of each data point in a cluster with their centroids. The red line (*k* = 3) indicates the number of clusters that has been designated. P-value of the nonparametric ANOVA test is reported inside the figures. * indicates p<0.05 ** p<0.01 and *** p<0.001 in the paired Wilcoxon tests between adjacent time points. Asterisks are reported only if p-values pass the Benjamini-Hochberg correction.

The three centroids for the four age groups are shown in Figure 4 A. We observe a gradual reduction in functional connectivity across all centroids as a function of age.

Dwell time of the first cluster shortened from 43.4 *±* 28 s to 12.4 *±* 8 s (ANOVA p=e-10); fractional occurrence from 0.46 *±* 0.1 to 0.23 *±* 0.1, (ANOVA p=e-9). In the third cluster, dwell time increased from 6.4 *±* 4 s to 19.3 *±* 9 s, (ANOVA p=e-15); fractional occurrence from 0.1 *±* 0.1 to 0.4 *±* 0.1 (ANOVA p=e-16).

On the other hand, the second cluster presented a decrease in dwell time (from 43.4 *±* 27 s to 9.7 *±* 4 s) and fractional occurrence (from 0.45 *±* 0.1 to 0.18 *±* 0.1) from to 3 to 11 months, and an increase of these measures (dwell time to 17.9 *±* 5 s and fractional occurrence to 0.37 *±* 0.1) from 11 to 17 months.

Spectral metastability of the first two clusters decreased with ageing, going from 3.2 ± 0.4 to 2.2 ± 0.4 (ANOVA p=e-11) for the first and from 3.0 ± 0.4 to 2.3 ± 0.6 for the second (ANOVA p=e-7). We did not observe a clear pattern of increase/decrease in spectral metastability of the third cluster.

### 3.3 Dynamic Functional Connectivity measures: Continuous EiDA

The position and speed plots for the **iPL** matrix in a representative rat are shown in Figure 5 A. Trajectories demonstrated significant changes with ageing. Patterns started as trajectories with a broad exploration of the configuration space and finished confined to a more compact area. Moreover, as measured below, the trajectories of younger rats exhibited a higher contribution from the first-eigenvector.

**Figure 5.**
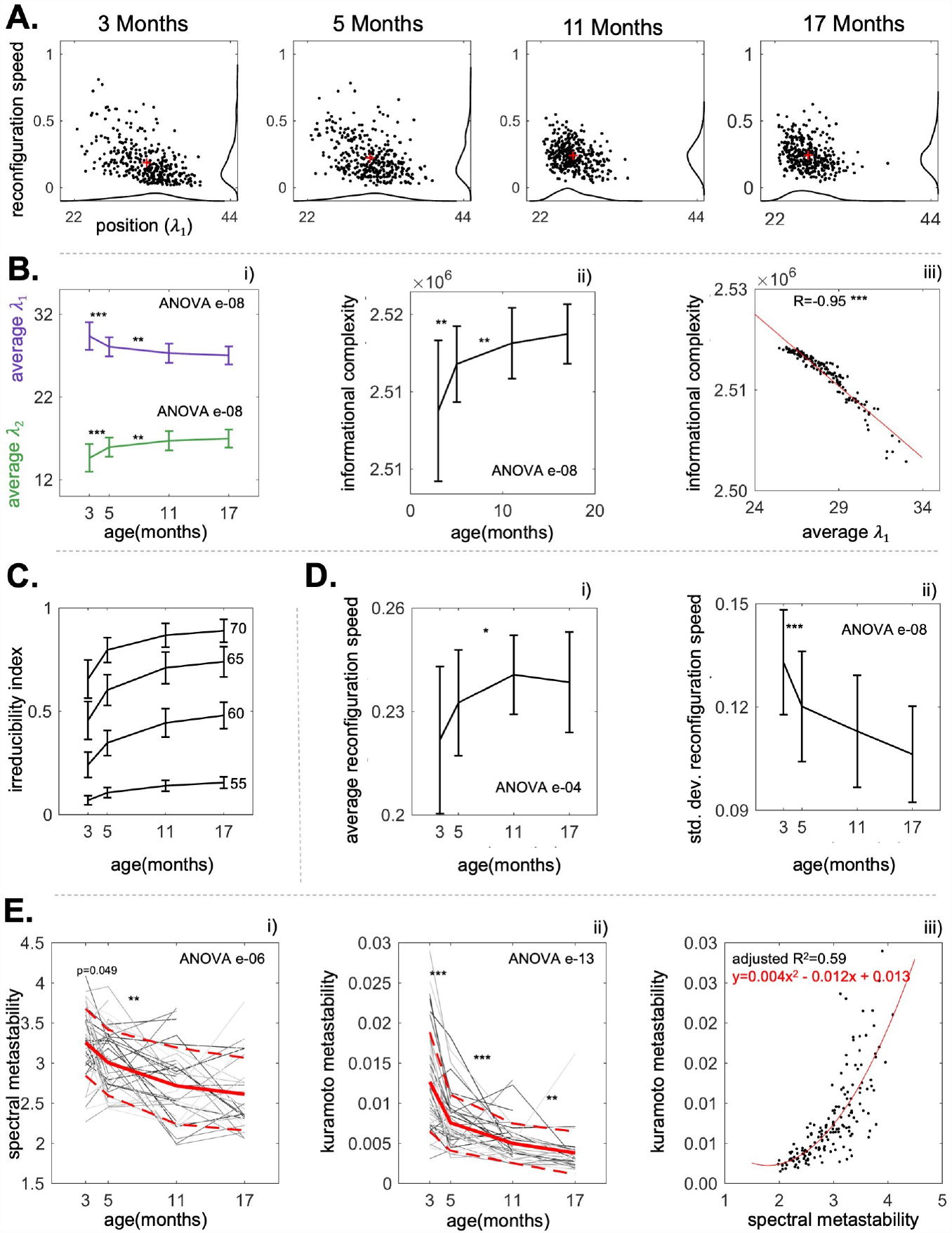
Results for the Continuous EiDA **A**. The position-speed evolution in a representative rat. The position is the first eigenvalue of the **iPL** matrix and the speed is the overall reconfiguration speed of the matrix (not considering eigenvectors separately). Red crosses represent the centers of mass of the distributions. Patterns evolve from a broader exploration of the space to a more compact one with age, where the first eigenvalue decreases and reconfiguration speeds grows. This means that connectivity patterns over time start from a more structured regime (less speed, higher eigenvalues) to a more random one. **B**. the measures of interest: (i) average eigenvalue per recording (mean ± variance, first eigenvalue in purple, second in green). The first eigenvalue shows a decrease over time, in line with the loss of structure that is noticeable in 6 B. (ii) informational complexity (iii) correlation between informational complexity and first eigenvalue, reflecting that the higher the first eigenvalue, the lower the amount of information contained in the iPL matrix. **C** Irreducibility Index, we plot 4 thresholds: 70, 65, 60, 55%. Plot has no significance indicated because we are not interested in the effect of ageing but on the fact that a non-negligible amount of the recordings is not irreducible to a single eigenvector. Moreover, for visual clarity, mean ± half of the variance is showed. **D**. (i) Average reconfiguration speed of the **iPL** matrix: it evolves faster with ageing (ii) standard deviation of the reconfiguration speed, quantifying the more compact exploration of the space associated with ageing. **E**. The evolution with ageing of (i) metastability and (ii) spectral metastability for all the *N* = 48 rats. Red lines indicate means, dotted lines indicate standard deviations. Here, the individual trajectories (gray lines) of each single rat are showed. Some lines present discontinuities in case the recording for a specific rat and age group was missing. Ageing is associated with a significant decrease of both metastability and our proposed measure of spectral metastability (iii) Scatter plot of spectral metastability versus metastability. The red line represents the quadratic fit that was performed to explain metastability as a function of spectral metastability. P-value of the nonparametric ANOVA test is reported inside the figures. * indicates p<0.05 ** p<0.01 and *** p<0.001 in the paired Wilcoxon tests between adjacent time points. Asterisks are reported only if p-values pass the Benjamini-Hochberg correction.

As shown in Figure 5 B.i, the average of the first eigenvalues per recording shows a statistically significant decay with ageing (from 29.5 ± 1.8 to 27.0 ± 1.0, ANOVA p=e-08). As in the case of static connectivity, this decay plateaued from 11 months to 17 months, suggesting that there is no significant change from middle-age to senescence. We observed an increase (ANOVA p=e-08) in the Informational Complexity measure defined in 2.9 with ageing that indicates that connectivity configurations evolve in a more random and less structured way in older rats compared with younger rats, as in Figure 5, B.ii. We correlated informational complexity with the average value of the first eigenvalue per recording, obtaining an overall correlation of -0.95, p<0.001 (Fig 5 B.iii). This establishes an information theoretical relation between the first eigenvalue and the amount of information contained in the **iPL** matrix, as also explained in Section 2.4.

Figure 5 C. shows the Irreducibility Index with four thresholds of 70, 65, 60 and 55%. The Irreducibility Index with a threshold of 65% is greater than 0.4 in 3 months rats and becomes greater than 0.7 in 17 months rats. This means that discarding the second eigenvector would neglect a significant (more than 35%) amount of functional connectivity information over the life-span of the rats. Same considerations hold for the Irreducibility Index with thresholds of 60 and 70 %. Information loss always increases with age.

We noticed an age-related increase in the average reconfiguration speed of **iPL** (Figure 5 D.i.) from 0.22 *±* 0.02 to 0.24 *±* 0.01 (ANOVA p=e-04) and a decrease in the standard deviation of the reconfiguration speed (Figure 5 D.ii.), from 0.13 *±* 0.02 to 0.11 *±* 0.01 (ANOVA p=e-08).

From the kinematic speed-displacement plots in Figure 5.A, we observed that as the rats age, there is a reduced exploration of the position-speed space. The decrease in the standard deviation of the reconfiguration speed quantifies this property. The second quantity is spectral metastability (standard deviation of “position”). We found that spectral metastability declines with age from 3.2 ± 0.4 to 2.6 ± 0.5 (ANOVA p=e-06) (Figure 5.E.i). Kuramoto metastability shows the same pattern of decay related to age. Indeed, spectral metastability and Kuramoto metastability are related: through a quadratic fit we found that the relation between spectral metastability (x) and the Kuramoto metastability (y) is the following: *y* = 0.004*x*^2^ − 0.012*x* + 0.013, adjusted R squared = 0.59 (Figure 5.E.iii). In conclusion, older brains are characterized by less variability both in the structure of the **iPL** matrix and in its rate of change.

We then applied Continuous EiDA to the two eigenvectors separately, gaining insight into the dynamic structural changes of the **iPL** matrix and its two building blocks. This offers a way to analyze brain network dynamics in a dimensionally reduced space, without any loss of information.

Figure 6 A. shows in the same representative rat of Figure 5 A., the 2-D embedding for the evolution of the two eigenvectors. Figure 6 B. shows the transformation from the raw data to the eigenvector evolution using EiDA, in a specific portion of the recording, for 3 months and 17 months. At 3 months signals are highly phase locked, as observable in the data. This is indicated by a more compact and ordered evolution of both eigenvectors: there is a clear dominant pattern of phase locking (first eigenvector, almost all brain areas in phase). On the other hand, in the 17 months case, both eigenvectors evolve more randomly and without a dominant underlying pattern.

**Figure 6.**
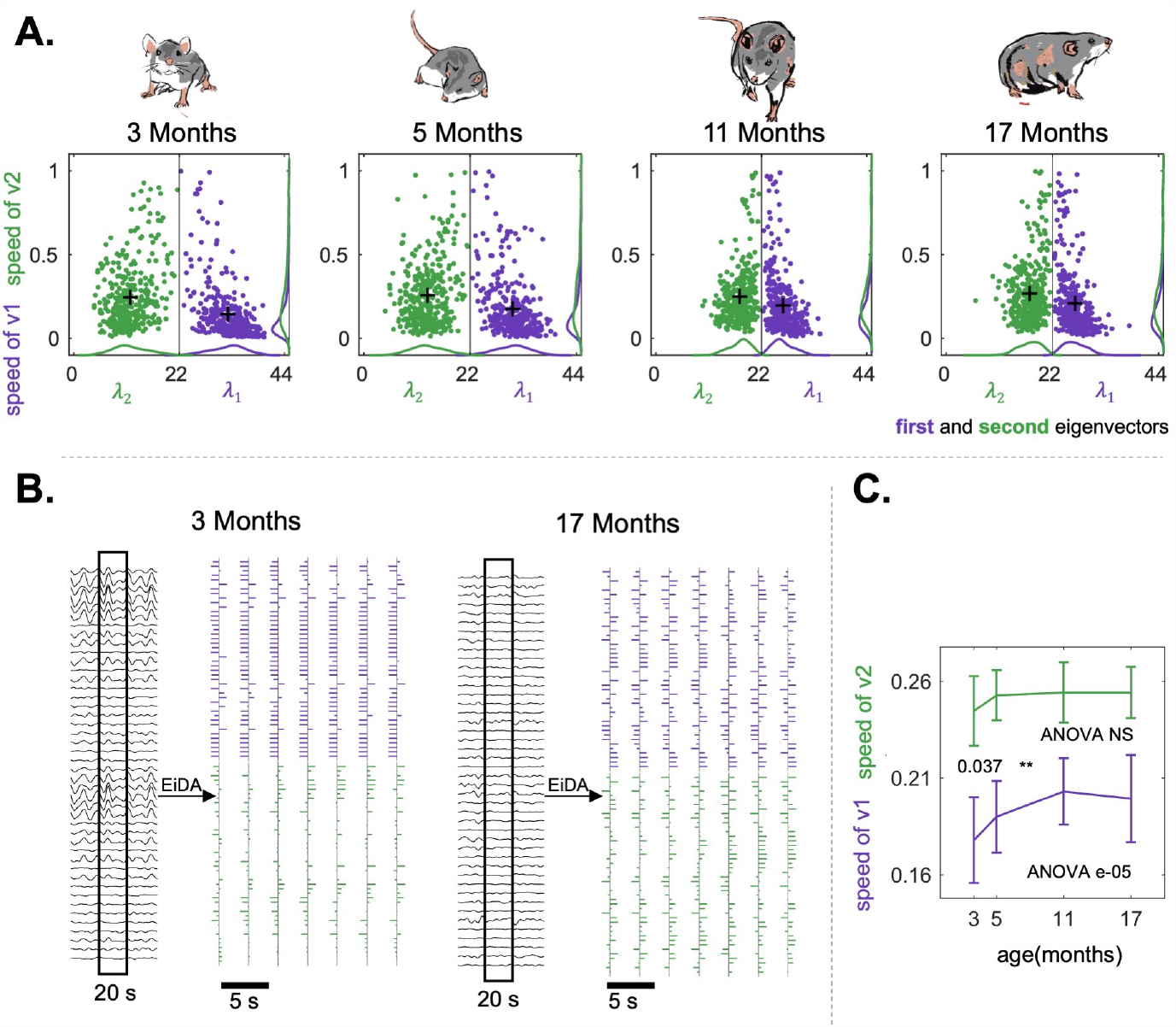
**A**. The position-speed plots of the two separate eigenvectors, for the same representative rat as in Figure 5 A. Here, the position for each eigenvectors is its associated eigenvalue, and the speed is the reconfiguration speed of the eigenvector. **B**. The EiDA transformation from the raw data to the evolution of the two eigenvectors for the same representative rat of **A**. in two different cases: 3 months (left) and 17 months (right). Note that in the first case we can observe higher instantaneous phase locking in the raw signals. This is related to a more ordered and compact evolution of eigenvectors, and the fact that the first eigenvector contains most of the information. In the second case, there is a lack of instantaneous phase locking. This is reflected by more random, less structured and equally informative eigenvectors. **C**. Average reconfiguration speed of first (purple) and second (green) eigenvector. We observe an increase in the speed of the first eigenvector. P-value of the nonparametric ANOVA test is reported inside the figures. * indicates p<0.05 ** p<0.01 and *** p<0.001 in the paired Wilcoxon tests between adjacent time points. Asterisks are reported only if p-values pass the Benjamini-Hochberg correction.

We observed (Figure 6 C.) a significant (ANOVA p= e-05) increase in the reconfiguration speed of the first eigenvector with ageing, from 0.18 ± 0.02 to 0.20 ± 0.02. Hence ageing is associated with a speed-up in the “walk” of the dominant pattern of connectivity. Notably, the evolution of the second eigenvector is always faster than the evolution of the first, as also visible in Figure 6 B. We also found that the reconfiguration speed of the overall matrix is strongly positively correlated with the reconfiguration speed of the first eigenvector (R=0.902, p<0.001), and positively correlated with the reconfiguration speed of the second eigenvector (R=0.6738, p<0.001).

### 3.4 Computation Times

We measured the time necessary to compute the EVD of the **iPL** matrix and compared our approach with the benchmark algorithm for LEiDA, implemented in Matlab, by varying the matrix size logarithmically from *N* = 10 to *N* = 10^5^ and generating 20 **iPL** matrices for each size. As shown in Figure 2 D, with our computer (Intel(R) Xeon(R) Silver 4116 CPU @ 2.10GHz), EiDA is 100 times faster for matrices with dimensions of 100×100, and becomes 1000 times faster for matrices larger than 10,000×10,000. Note that the matrix decomposition in the two eigenvectors also allows to save each matrix with 2*N* elements instead of 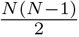, offering a significant advantage in terms of RAM requirements. Computational speed should ease the extension of EiDA to higher dimensionality both in space and time.

## 4 Discussion

In this study we explore the implications of using phase relationships and the **iPL**, which we have analytically characterized. We investigate reconfiguration dynamics in addition to state dynamics, and we test our novel adapted method on the resting-state fMRI data from a study of ageing in rats. Our resulting method, EiDA, provides an analytically based framework for dimensionality reduction and dynamical analysis. Here we showcase its utility from a number of theoretical and application perspectives.

### 4.1 Eigenvector decomposition of the iPL matrix

EiDA provides an analytical solution expressible in closed form for the lossless decomposition of the instantaneous Phase Locking matrix. Complete information is encoded in two eigenvectors and one eigenvalue, allowing the calculation of conventional dynamic metrics and the derivation of novel measures to further characterize the dynamic evolution of the spontaneous activity in the resting-state brain. Additionally, the analytical methods significantly accelerate the calculation of the orthogonal decomposition and enable its extension to much higher dimensional data, either or both in space and time.

### 4.2 Role of the eigenvalues and eigenvectors of the iPL matrix

As EiDA is an eigenvector decomposition of the **iPL** matrix, one could be tempted to interpret the eigenspace spanned by first eigenvector as the denoised form of the matrix. However this may not be necessarily true as there is no clear differentiation of signal and noise, particularly in the regime where *λ*_1_ is far from *N* (its maximum possible value). One could articulate an interpretation of the leading eigenvector as the container of the large scale information about the connectivity patterns, while the second eigenvector contains more localized details, as in Figure 6 B.i. and Figure 2 C.i., following ideas from graph signal analysis (63). This interpretation is possible only if the first eigenvalue is significantly higher than the second. This interpretation breaks down in the regime where *λ*_1_ ≈ *λ*_2_ as shown by the counter example of Figure 2 B., where in a fully structured matrix, made of 4 blocks, the two eigenvalues are equal, and using only the first eigenvector would discard 50% of the overall phase locking information. In conclusion, both eigenvectors and eigenvalues are necessary to interpret the **iPL** matrix.

The analytical derivation in this paper allows the rigorous definition of a global parameter summarising the heterogeneity in the signal phases, the first eigenvalue or spectral radius of the **iPL** matrix, whose distribution over time contains the information both on global connectivity and on informational complexity of the matrix; the standard deviation of this distribution we have defined as a novel estimator of metastability.

This eigenvalue can be a more accurate indicator than the Kuramoto Order Parameter of the phase locking structure, being the 2-norm of the phase locking matrix, rather than the center of mass of instantaneous angles. For example, in a maximally synchronized and ordered state, as in Figure 2 A., where all the signals are either in phase or in antiphase, the Kuramoto Order Parameter can be 0 if half of the signals are in phase and the other half in antiphase, whilst the first eigenvector would reach its maximal possible value and capture the “singular” structure of this configuration.

### 4.3 Discrete and Continuous EiDA

Discrete EiDA can be used to extract discrete brain modes whenever the time-series of the two eigenvectors would warrant the use of clustering; and in addition, Continuous EiDA can provide a complementary perspective to discrete states by capturing the evolution of the dynamics in 2-dimensional plots. Continuous EiDA can also be applied within the Discrete EiDA states. Metrics from both discrete and Continuous EiDA may provide more accurate estimations of differences between case and control groups, and so may improve discrimination and classification accuracy of putative neuromechanistic biomarkers (23).

### 4.4 Dynamical and complexity insights for ageing

The results of static analysis show a decrease of connectivity through ageing. The dynamic analysis corroborates and expands on this, grounding the results obtained with the dynamic approach. Where previous results ((58)) were limited to the default mode network, here we are considering the dynamic of the brain as a whole.

#### 4.4.1 Discrete EiDA findings

Discrete EiDA captured the changes in spatiotemporal patterns of connectivity associated with age. These patterns showed a loss of phase synchrony and a reduced connectivity structure with ageing. This likely explains the increase in fractional occurrence and dwell times of the third centroids, that are associated with disrupted brain synchrony, and the decrease of fractional occurrence and dwell times of the first two centroids, which relate to phase-locked states. This is also in line with previous results from ageing studies obtained using different methodologies, exploring both static and dynamic functional connectivity ((36; 32; 38; 39; 29; 64)). This increase in randomness is also shown by the fact that the second centroid loses its structure at the age of 17 months (Figure 4 A). This means that at the age of 17 months there is a rearrangement of centroids when connectivity breaks down with ageing. This explains also why fractional occurrence and dwell time of the second cluster increase again at the age of 17 months, after having decreased.

#### 4.4.2 Continous EiDA findings

Our interpretable (“position” = overall connectivity and “speed”) 2-D embedding allowed to visualize the effect of ageing on dynamic Functional Connectivity. Older brains are less flexible (less variability) in the exploration of connectivity patterns and in the speed of the exploration: the “filling” of the space becomes more compact with age. Both Continuous EiDA applied to the full **iPL** matrix and to the separate eigenvectors, with their associated measures, highlighted this result. Indeed, a decrease in spectral metastability, both in the overall recordings and in the separate EiDA clusters, was observed. Metastability is related to cognitive flexibility and adaptability to the environment ((65)), and the decrease we found is in line with previous findings, where a proxy measure of metastability was computed based on spatial diversity ((64)).

The increase of the reconfiguration speed of the first eigenvector associated with ageing is also intriguing as the brain dynamics is shown to move from a relatively coherent exploration of the kinematic space to a more random exploration with ageing, as pointed out by (35).

Continuous EiDA showed also an increase of the dynamic complexity of dFC. Increases in complexity with healthy ageing have previously been demonstrated using point-wise correlation dimension (66) and multi-scale entropy (67; 68) in resting-state neuroimaging which is congruent with our findings. Taken together, our findings are particularly compelling as similar results are obtained using two distinct methods grounded in different theoretical frameworks: dynamical system analysis and information theory. Furthermore, by using our analytical interpretation and by correlating eigenvalues with informational complexity, we have established a link between dynamic connectivity analysis and the notion of informational complexity of signals.

Retaining only the leading eigenvector (as in LEiDA), appears to result in age-related increase in dFC information loss. This implies that, in ageing, more information is encoded in the 2nd eigenvector. This is interesting as one can thus interpret the 2nd eigenvector to be more related to the localized connectivity information (63). Accordingly, previous studies (67; 68) have shown that healthy ageing is associated with a shift in the local vs global balance, with less information coded in global interactions and more in the local dynamics.

Utilising this methodology we discovered multiple sensitive markers of the ageing brain:

- The average position, the first eigenvalue of the iPL matrix, which is related to the total quota of connectivity and the informational complexity of the **iPL** matrix.
- The dwell time and fractional occurrence of the brain states, which yield the highest significance (e-10,e-10 for the first state; e-15,e-12 for the second; e-15,e-16 for the third) and which inform about the dynamic composition of the changes in connectivity patterns.
- A newly introduced measure, spectral metastability, which can be computed both in the overall recording and in the separate Discrete EiDA modes. This measure is more general than Kuramoto metastability because it can be calculated from any time varying matrix, for example a sliding window correlation matrix (as it is just the standard deviation in time of the first eigenvalue). Moreover, spectral metastability takes into account the full structure of the **iPL** matrix, while the Kuramoto metastability uses the center of mass of the phases. Therefore, spectral metastability is superior to Kuramoto for the analysis of variability of the full **iPL** pattern (see Sections 2.5 and 2.6). Metastability has proven to be a candidate biomarker of schizophrenia pathology ((23)) and has also been linked also to neurodevelopmental changes ((22)).

These newly introduced measures are not only more sensitive than static measures but also provide deeper insights into the dynamics of the brain.

### 4.5 Conclusion

EiDA provides a computationally fast, analytically based, closed form method to extract connectivity information in a lossless manner from signals in high-dimensional time-series. Its application to fMRI data from a longitudinal rat ageing study demonstrated its advantages over conventional methods and revealed a strong relationship between dynamical and informational metrics. This approach holds great promise in enhancing the accuracy of potential neuromechanistic biomarkers for diseases or the assessment of intervention studies. Furthermore, its potential extends beyond neuroscience, opening doors to various applications.

Its high speed and memory efficiency paves the way for advancements that would have been computationally unfeasible so far, like: a) Real-time algorithms ((69; 70; 71)) b) Studies with a high number of brain signals, where parcellation is extremely fine-grained or absent.

In summary, with this work we have introduced

- A full characterization of the **iPL** matrix and its properties.
- An analytical and therefore ultra-fast way of decomposing it.
- A new algorithm (Discrete EiDA) for the clustering of brain states, that can ease RAM requirements by orders of magnitude. Moreover, Discrete EiDA solves the “flipping sign” problem that was observed in LEiDA (see Section 2.8.1).
- A 2-d embedding of dFC and two new measures for the continuous analysis of the connectivity patterns.

We emphasise that all these concepts may be extended to any square symmetric dFC matrix, after the choice of an adequate number of eigenvectors, which may not necessarily be two, making EiDA a general and comprehensive approach for dynamic Functional connectivity.

The code for EiDA is available at www.github.com/alteriis.

## 5 Appendix

### 5.1 Analytical Computation of the Eigenvectors of iPL Matrix

Given that the matrix **iPL**_*ij*_ = cos(*θ*_*i*_ − *θ*_*j*_), and given that cos(*x* − *y*) = cos(*x*) cos(*y*) + sin(*x*) sin(*y*), then **iPL**_*ij*_ = cos(*θ*_*i*_) cos(*θ*_*j*_) + sin(*θ*_*i*_) sin(*θ*_*j*_). This means that the *iPL* matrix can be decomposed in the sum of two matrices. We define the two vectors, the “cosine” vector **c** = (cos(*θ*_1_), cos(*θ*_2_), …, cos(*θ*_*N*_))^*T*^ ∈ ℝ^*N*^, and the “sine” vector **s** = (sin(*θ*_1_), sin(*θ*_2_), …, sin(*θ*_*N*_))^*T*^ ∈ ℝ^*N*^, and rewrite the matrix as **iPL** = **cc**^*T*^ + **ss**^*T*^.

This matrix is symmetric and has ones on the diagonal, as **iPL**_*ii*_ = cos(*θ*_*i*_ − *θ*_*i*_) = cos(0) = 1 Its trace is thus *Tr*(**iPL**) = *N*. Importantly, this decomposition demonstrates that the matrix is a rank 2 matrix, which, being symmetric and positive semidefinite, will only have 2 non null (and greater than zero) eigenvalues *λ*_1_ and *λ*_2_ and their associated eigenvectors (as observed by (21)). This also implies that, as the sum of the eigenvalues of a matrix is equal to the trace of the matrix, *Tr*(**iPL**) = *N* = *λ*_1_ + *λ*_2_. Moreover, the two non-trivial eigenvectors will be a linear combination of **c** and **s**, so they will be of the form *A***c** + *B***s**. Thus, we only need to compute the two scalar values *A* and *B* to find the eigenvectors. The eigenvalue equation **iPL**(*A***c** + *B***s**) = *λ*(*A***c** + *B***s**) means that:

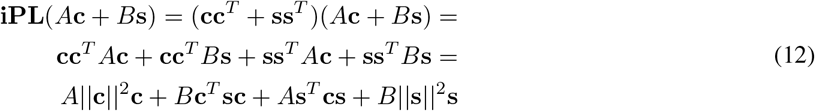

Let us define the following quantities:

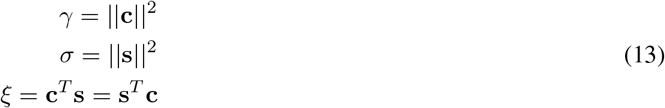

We can then rewrite **iPL**(*A***c** + *B***s**) = (*Aγ* + *Bξ*)**c** + (*Aξ* + *Bσ*)**s**, which needs to be equal to *λ*(*A***c** + *B***s**). We therefore have a system of equations:

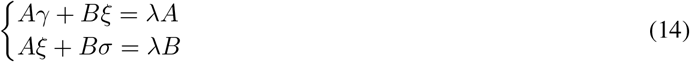

By dividing the two equations we obtain: 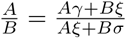. Since the eigenvectors remain eigenvectors if they are divided by an arbitrary constant, we impose that *A* = 1. Therefore, we obtain *B*^2^*ξ* + *B*(*γ σ*) *ξ* = 0, so, Δ = (*γ − σ*)^2^ + 4*ξ*^2^ and:

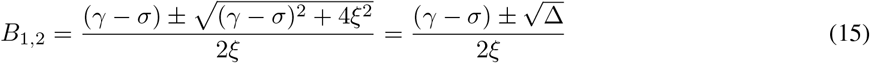

So, the two eigenvectors **v**_**1**_ and **v**_**2**_ are:

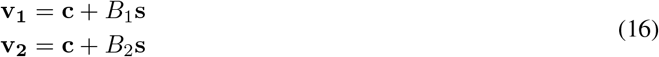

### 5.2 Analytical computation of Eigenvalues of iPL Matrix

To compute the eigenvalues we have to show first that 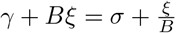 for both *B*_1_ and *B*_2_. We prove it for *B*_1_, the proof for *B*_2_ is analogous:

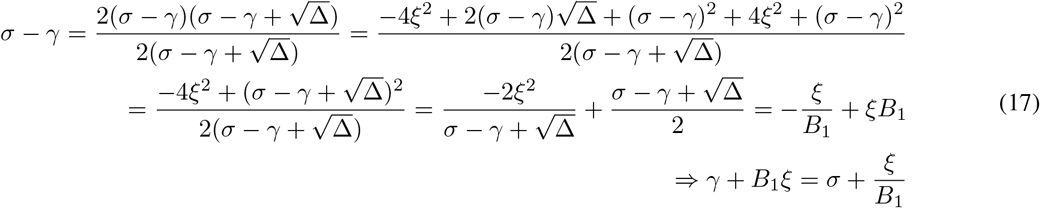

Using again the eigenvalue equation, we compute the eigenvalue *λ*_1_:

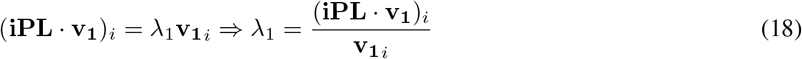

where

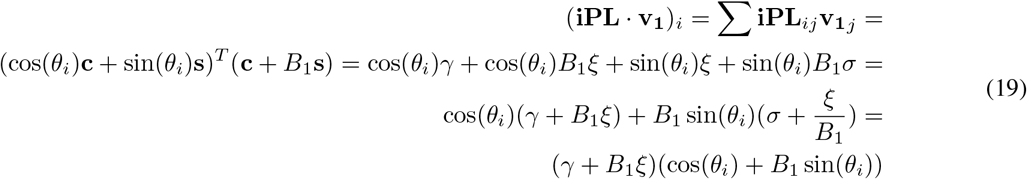

So, given that **v**_**1***i*_ = cos(*θ*_*i*_) + *B*_1_ sin(*θ*_*i*_), we have that:

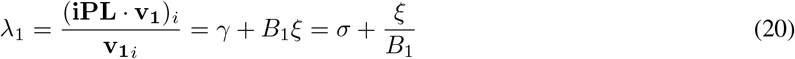

and we obtain *λ*_2_ similarly:

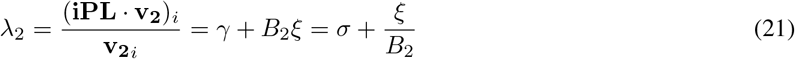

