## Supplementary material for "EiDA: A lossless approach for the dynamic analysis of connectivity patterns in signals; application to resting state fMRI of a model of ageing": supplementary_material_statistics.docx

```
FIGURE 3

measure: FC index for ipl matrix
Nonparametric ANOVA (kruskal wallis) p-value: 9.330767e-09

      Source             SS            df             MS             Chi-sq        Prob>Chi-sq
    ___________    ______________    _______    ______________    ____________    ______________

    {'Columns'}    {[4.8729e+04]}    {[  3]}    {[1.6243e+04]}    {[ 40.2719]}    {[9.3308e-09]}
    {'Error'  }    {[9.5261e+04]}    {[116]}    {[  821.2155]}    {0×0 double}    {0×0 double  }
    {'Total'  }    {[    143990]}    {[119]}    {0×0 double  }    {0×0 double}    {0×0 double  }


T1 vs T2  wilcoxon p value = 0.00089443
        ---------
T2 vs T3  wilcoxon p value = 0.007271
        ---------
T3 vs T4  wilcoxon p value = 0.40483
        ---------

means:1.743940e+01 1.432518e+01 1.238604e+01 1.202920e+01
stds:3.783279e+00 2.822229e+00 2.334583e+00 2.665591e+00

                              ~~~~~~~~~~~~~~~~~~~~~~~~~~~~~~~~~~~

measure: FC index correlation matrix
Nonparametric ANOVA (kruskal wallis) p-value: 9.854890e-09

      Source             SS            df             MS             Chi-sq        Prob>Chi-sq
    ___________    ______________    _______    ______________    ____________    ______________

    {'Columns'}    {[4.8594e+04]}    {[  3]}    {[1.6198e+04]}    {[ 40.1599]}    {[9.8549e-09]}
    {'Error'  }    {[9.5396e+04]}    {[116]}    {[  822.3833]}    {0×0 double}    {0×0 double  }
    {'Total'  }    {[    143990]}    {[119]}    {0×0 double  }    {0×0 double}    {0×0 double  }


T1 vs T2  wilcoxon p value = 0.0010357
        ---------
T2 vs T3  wilcoxon p value = 0.0025846
        ---------
T3 vs T4  wilcoxon p value = 0.50383
        ---------

means:1.969628e+01 1.678065e+01 1.408462e+01 1.367809e+01
stds:3.878135e+00 3.350128e+00 2.939263e+00 3.123053e+00


FIGURE 4

measure: fractional occurrence of cluster 1
Nonparametric ANOVA (kruskal wallis) p-value: 6.760030e-10

      Source             SS            df             MS             Chi-sq        Prob>Chi-sq
    ___________    ______________    _______    ______________    ____________    ______________

    {'Columns'}    {[5.5215e+04]}    {[  3]}    {[1.8405e+04]}    {[ 45.6413]}    {[6.7600e-10]}
    {'Error'  }    {[8.8745e+04]}    {[116]}    {[  765.0473]}    {0×0 double}    {0×0 double  }
    {'Total'  }    {[    143960]}    {[119]}    {0×0 double  }    {0×0 double}    {0×0 double  }


T1 vs T2  wilcoxon p value = 0.31844
        ---------
T2 vs T3  wilcoxon p value = 0.73428
        ---------
T3 vs T4  wilcoxon p value = 1.6378e-05
        ---------

means:4.552682e-01 4.352490e-01 4.273946e-01 2.345785e-01
stds:1.079040e-01 9.034125e-02 8.729963e-02 1.137359e-01

                              ~~~~~~~~~~~~~~~~~~~~~~~~~~~~~~~~~~~

measure: avg duration of cluster 1
Nonparametric ANOVA (kruskal wallis) p-value: 1.487152e-10

      Source             SS            df             MS             Chi-sq        Prob>Chi-sq
    ___________    ______________    _______    ______________    ____________    ______________

    {'Columns'}    {[5.8966e+04]}    {[  3]}    {[1.9655e+04]}    {[ 48.7326]}    {[1.4872e-10]}
    {'Error'  }    {[8.5023e+04]}    {[116]}    {[  732.9542]}    {0×0 double}    {0×0 double  }
    {'Total'  }    {[1.4399e+05]}    {[119]}    {0×0 double  }    {0×0 double}    {0×0 double  }


T1 vs T2  wilcoxon p value = 0.019569
        ---------
T2 vs T3  wilcoxon p value = 0.075213
        ---------
T3 vs T4  wilcoxon p value = 0.00014773
        ---------

means:4.339735e+01 2.676665e+01 2.234757e+01 1.240051e+01
stds:2.824505e+01 1.112871e+01 5.111319e+00 8.387121e+00

                              ~~~~~~~~~~~~~~~~~~~~~~~~~~~~~~~~~~~

measure: metastability of cluster 1
Nonparametric ANOVA (kruskal wallis) p-value: 2.185427e-11

      Source           SS          df             MS             Chi-sq        Prob>Chi-sq
    ___________    __________    _______    ______________    ____________    ______________

    {'Columns'}    {[ 63697]}    {[  3]}    {[2.1232e+04]}    {[ 52.6421]}    {[2.1854e-11]}
    {'Error'  }    {[ 80293]}    {[116]}    {[  692.1810]}    {0×0 double}    {0×0 double  }
    {'Total'  }    {[143990]}    {[119]}    {0×0 double  }    {0×0 double}    {0×0 double  }


T1 vs T2  wilcoxon p value = 0.46528
        ---------
T2 vs T3  wilcoxon p value = 0.0027653
        ---------
T3 vs T4  wilcoxon p value = 5.307e-05
        ---------

means:3.182195e+00 3.063381e+00 2.734559e+00 2.189846e+00
stds:4.141962e-01 4.858451e-01 4.569723e-01 3.509641e-01

                              ~~~~~~~~~~~~~~~~~~~~~~~~~~~~~~~~~~~

measure: fractional occurrence of cluster 2
Nonparametric ANOVA (kruskal wallis) p-value: 1.173425e-12

      Source             SS            df             MS             Chi-sq        Prob>Chi-sq
    ___________    ______________    _______    ______________    ____________    ______________

    {'Columns'}    {[7.0884e+04]}    {[  3]}    {[2.3628e+04]}    {[ 58.5945]}    {[1.1734e-12]}
    {'Error'  }    {[7.3075e+04]}    {[116]}    {[  629.9579]}    {0×0 double}    {0×0 double  }
    {'Total'  }    {[1.4396e+05]}    {[119]}    {0×0 double  }    {0×0 double}    {0×0 double  }


T1 vs T2  wilcoxon p value = 0.082182
        ---------
T2 vs T3  wilcoxon p value = 4.2782e-06
        ---------
T3 vs T4  wilcoxon p value = 4.2807e-06
        ---------

means:4.518199e-01 4.117816e-01 1.847701e-01 3.698276e-01
stds:1.056049e-01 1.018370e-01 1.016747e-01 6.536700e-02

                              ~~~~~~~~~~~~~~~~~~~~~~~~~~~~~~~~~~~

measure: avg duration of cluster 2
Nonparametric ANOVA (kruskal wallis) p-value: 1.477730e-15

      Source             SS            df             MS             Chi-sq        Prob>Chi-sq
    ___________    ______________    _______    ______________    ____________    ______________

    {'Columns'}    {[8.7301e+04]}    {[  3]}    {[2.9100e+04]}    {[ 72.1509]}    {[1.4777e-15]}
    {'Error'  }    {[5.6686e+04]}    {[116]}    {[  488.6763]}    {0×0 double}    {0×0 double  }
    {'Total'  }    {[1.4399e+05]}    {[119]}    {0×0 double  }    {0×0 double}    {0×0 double  }


T1 vs T2  wilcoxon p value = 0.0033789
        ---------
T2 vs T3  wilcoxon p value = 1.9209e-06
        ---------
T3 vs T4  wilcoxon p value = 2.3704e-05
        ---------

means:4.339217e+01 2.658974e+01 9.669624e+00 1.791753e+01
stds:2.655078e+01 1.339043e+01 4.336543e+00 5.271858e+00

                              ~~~~~~~~~~~~~~~~~~~~~~~~~~~~~~~~~~~

measure: metastability of cluster 2
Nonparametric ANOVA (kruskal wallis) p-value: 4.421238e-07

      Source             SS            df             MS             Chi-sq        Prob>Chi-sq
    ___________    ______________    _______    ______________    ____________    ______________

    {'Columns'}    {[3.9140e+04]}    {[  3]}    {[1.3047e+04]}    {[ 32.3475]}    {[4.4212e-07]}
    {'Error'  }    {[1.0485e+05]}    {[116]}    {[  903.8753]}    {0×0 double}    {0×0 double  }
    {'Total'  }    {[    143990]}    {[119]}    {0×0 double  }    {0×0 double}    {0×0 double  }


T1 vs T2  wilcoxon p value = 0.027029
        ---------
T2 vs T3  wilcoxon p value = 0.00011499
        ---------
T3 vs T4  wilcoxon p value = 0.50383
        ---------

means:2.957670e+00 2.753671e+00 2.376982e+00 2.295182e+00
stds:3.843105e-01 4.517109e-01 4.039355e-01 5.554944e-01

                              ~~~~~~~~~~~~~~~~~~~~~~~~~~~~~~~~~~~

measure: fractional occurrence of cluster 3
Nonparametric ANOVA (kruskal wallis) p-value: 1.236732e-16

      Source             SS            df             MS             Chi-sq        Prob>Chi-sq
    ___________    ______________    _______    ______________    ____________    ______________

    {'Columns'}    {[9.3374e+04]}    {[  3]}    {[3.1125e+04]}    {[ 77.1777]}    {[1.2367e-16]}
    {'Error'  }    {[5.0599e+04]}    {[116]}    {[  436.1980]}    {0×0 double}    {0×0 double  }
    {'Total'  }    {[    143973]}    {[119]}    {0×0 double  }    {0×0 double}    {0×0 double  }


T1 vs T2  wilcoxon p value = 0.0064144
        ---------
T2 vs T3  wilcoxon p value = 4.072e-06
        ---------
T3 vs T4  wilcoxon p value = 0.4112
        ---------

means:9.291188e-02 1.529693e-01 3.878352e-01 3.955939e-01
stds:9.955840e-02 9.829296e-02 1.224521e-01 9.764673e-02
measure: avg duration of cluster 3
Nonparametric ANOVA (kruskal wallis) p-value: 2.323208e-15

      Source             SS            df             MS             Chi-sq        Prob>Chi-sq
    ___________    ______________    _______    ______________    ____________    ______________

    {'Columns'}    {[8.6181e+04]}    {[  3]}    {[2.8727e+04]}    {[ 71.2335]}    {[2.3232e-15]}
    {'Error'  }    {[5.7789e+04]}    {[116]}    {[  498.1843]}    {0×0 double}    {0×0 double  }
    {'Total'  }    {[    143970]}    {[119]}    {0×0 double  }    {0×0 double}    {0×0 double  }


T1 vs T2  wilcoxon p value = 0.004114
        ---------
T2 vs T3  wilcoxon p value = 3.7243e-05
        ---------
T3 vs T4  wilcoxon p value = 0.57165
        ---------

means:6.379724e+00 9.145384e+00 1.904508e+01 1.925484e+01
stds:4.153914e+00 4.676275e+00 7.363499e+00 8.649783e+00

                              ~~~~~~~~~~~~~~~~~~~~~~~~~~~~~~~~~~~

measure: metastability of cluster 3
Nonparametric ANOVA (kruskal wallis) p-value: 5.250033e-04

      Source             SS            df             MS             Chi-sq        Prob>Chi-sq
    ___________    ______________    _______    ______________    ____________    ______________

    {'Columns'}    {[1.9936e+04]}    {[  3]}    {[6.6454e+03]}    {[ 17.6272]}    {[5.2500e-04]}
    {'Error'  }    {[1.1013e+05]}    {[112]}    {[  983.2919]}    {0×0 double}    {0×0 double  }
    {'Total'  }    {[    130065]}    {[115]}    {0×0 double  }    {0×0 double}    {0×0 double  }


T1 vs T2  wilcoxon p value = 0.16973
        ---------
T2 vs T3  wilcoxon p value = 0.021286
        ---------
T3 vs T4  wilcoxon p value = 0.00020859
        ---------

means:2.329107e+00 2.482088e+00 2.202171e+00 2.648467e+00
stds:5.089839e-01 4.951227e-01 3.080878e-01 3.898462e-01


FIGURE 5

measure: eigenvalue 1 = spectral radius
Nonparametric ANOVA (kruskal wallis) p-value: 1.030758e-08

      Source             SS            df             MS             Chi-sq        Prob>Chi-sq
    ___________    ______________    _______    ______________    ____________    ______________

    {'Columns'}    {[4.8482e+04]}    {[  3]}    {[1.6161e+04]}    {[ 40.0679]}    {[1.0308e-08]}
    {'Error'  }    {[9.5508e+04]}    {[116]}    {[  823.3431]}    {0×0 double}    {0×0 double  }
    {'Total'  }    {[    143990]}    {[119]}    {0×0 double  }    {0×0 double}    {0×0 double  }


T1 vs T2  wilcoxon p value = 0.00089443
        ---------
T2 vs T3  wilcoxon p value = 0.0038542
        ---------
T3 vs T4  wilcoxon p value = 0.54401
        ---------

means:2.946121e+01 2.805408e+01 2.713789e+01 2.702104e+01
stds:1.835784e+00 1.276381e+00 1.005053e+00 1.076974e+00

                              ~~~~~~~~~~~~~~~~~~~~~~~~~~~~~~~~~~~

measure: informational complexity
Nonparametric ANOVA (kruskal wallis) p-value: 1.250180e-08

      Source             SS            df             MS             Chi-sq        Prob>Chi-sq
    ___________    ______________    _______    ______________    ____________    ______________

    {'Columns'}    {[4.8004e+04]}    {[  3]}    {[1.6001e+04]}    {[ 39.6725]}    {[1.2502e-08]}
    {'Error'  }    {[9.5986e+04]}    {[116]}    {[  827.4651]}    {0×0 double}    {0×0 double  }
    {'Total'  }    {[1.4399e+05]}    {[119]}    {0×0 double  }    {0×0 double}    {0×0 double  }


T1 vs T2  wilcoxon p value = 0.0012866
        ---------
T2 vs T3  wilcoxon p value = 0.0038542
        ---------
T3 vs T4  wilcoxon p value = 0.31848
        ---------

means:2.509426e+06 2512707 2.514419e+06 2.514659e+06
stds:4.898115e+03 2.640916e+03 1.793370e+03 1.891798e+03


     				 ~~~~~~~~~~~~~~~~~~~~~~~~~~~~~~~~~~~

measure: std dev of reconfiguration speed
Nonparametric ANOVA (kruskal wallis) p-value: 4.638456e-08

      Source             SS            df             MS             Chi-sq        Prob>Chi-sq
    ___________    ______________    _______    ______________    ____________    ______________

    {'Columns'}    {[4.4750e+04]}    {[  3]}    {[1.4917e+04]}    {[ 36.9835]}    {[4.6385e-08]}
    {'Error'  }    {[9.9240e+04]}    {[116]}    {[  855.5167]}    {0×0 double}    {0×0 double  }
    {'Total'  }    {[    143990]}    {[119]}    {0×0 double  }    {0×0 double}    {0×0 double  }


T1 vs T2  wilcoxon p value = 0.00061564
        ---------
T2 vs T3  wilcoxon p value = 0.062683
        ---------
T3 vs T4  wilcoxon p value = 0.14139
        ---------

means:1.333286e-01 1.188397e-01 1.115129e-01 1.062699e-01
stds:1.524897e-02 1.618577e-02 1.536319e-02 1.397473e-02

                              ~~~~~~~~~~~~~~~~~~~~~~~~~~~~~~~~~~~

measure: mean reconfiguration speed
Nonparametric ANOVA (kruskal wallis) p-value: 1.669891e-04

      Source             SS            df             MS             Chi-sq        Prob>Chi-sq
    ___________    ______________    _______    ______________    ____________    ______________

    {'Columns'}    {[2.4241e+04]}    {[  3]}    {[8.0805e+03]}    {[ 20.0343]}    {[1.6699e-04]}
    {'Error'  }    {[1.1975e+05]}    {[116]}    {[1.0323e+03]}    {0×0 double}    {0×0 double  }
    {'Total'  }    {[    143990]}    {[119]}    {0×0 double  }    {0×0 double}    {0×0 double  }


T1 vs T2  wilcoxon p value = 0.10201
        ---------
T2 vs T3  wilcoxon p value = 0.013975
        ---------
T3 vs T4  wilcoxon p value = 0.51705
        ---------

means:2.212638e-01 2.321094e-01 2.424817e-01 2.384687e-01
stds:2.139190e-02 1.632248e-02 1.028617e-02 1.455955e-02


~~~~~~~~~~~~~~~~~~~~~~~~~~~~~~~~~~~

measure: spectral metastability
Nonparametric ANOVA (kruskal wallis) p-value: 7.790465e-07

      Source             SS            df             MS             Chi-sq        Prob>Chi-sq
    ___________    ______________    _______    ______________    ____________    ______________

    {'Columns'}    {[3.7728e+04]}    {[  3]}    {[1.2576e+04]}    {[ 31.1799]}    {[7.7905e-07]}
    {'Error'  }    {[1.0626e+05]}    {[116]}    {[  916.0546]}    {0×0 double}    {0×0 double  }
    {'Total'  }    {[    143990]}    {[119]}    {0×0 double  }    {0×0 double}    {0×0 double  }


T1 vs T2  wilcoxon p value = 0.049498
        ---------
T2 vs T3  wilcoxon p value = 0.0012866
        ---------
T3 vs T4  wilcoxon p value = 0.76552
        ---------

means:3.236936e+00 3.005686e+00 2.653202e+00 2.614077e+00
stds:4.379134e-01 4.394796e-01 4.375050e-01 4.519052e-01

                              ~~~~~~~~~~~~~~~~~~~~~~~~~~~~~~~~~~~

measure: kuramoto metastability
Nonparametric ANOVA (kruskal wallis) p-value: 4.437007e-13

      Source             SS            df             MS             Chi-sq        Prob>Chi-sq
    ___________    ______________    _______    ______________    ____________    ______________

    {'Columns'}    {[7.3292e+04]}    {[  3]}    {[2.4431e+04]}    {[ 60.5717]}    {[4.4370e-13]}
    {'Error'  }    {[7.0698e+04]}    {[116]}    {[  609.4672]}    {0×0 double}    {0×0 double  }
    {'Total'  }    {[    143990]}    {[119]}    {0×0 double  }    {0×0 double}    {0×0 double  }


T1 vs T2  wilcoxon p value = 5.7924e-05
        ---------
T2 vs T3  wilcoxon p value = 0.00028308
        ---------
T3 vs T4  wilcoxon p value = 0.0031618
        ---------

means:1.303540e-02 7.771787e-03 4.630907e-03 3.795565e-03
stds:6.643573e-03 4.018835e-03 2.155167e-03 2.661123e-03


FIGURE 6
```

measure: reconfiguration speed 1^st^ eigenvector

```
Nonparametric ANOVA (kruskal wallis) p-value: 1.366842e-05

      Source             SS            df             MS             Chi-sq        Prob>Chi-sq
    ___________    ______________    _______    ______________    ____________    ______________

    {'Columns'}    {[3.0556e+04]}    {[  3]}    {[1.0185e+04]}    {[ 25.2532]}    {[1.3668e-05]}
    {'Error'  }    {[1.1343e+05]}    {[116]}    {[  977.8764]}    {0×0 double}    {0×0 double  }
    {'Total'  }    {[    143990]}    {[119]}    {0×0 double  }    {0×0 double}    {0×0 double  }


T1 vs T2  wilcoxon p value = 0.036826
        ---------
T2 vs T3  wilcoxon p value = 0.0053197
        ---------
T3 vs T4  wilcoxon p value = 0.23694
        ---------

means:1.784098e-01 1.903294e-01 2.061423e-01 1.996177e-01
stds:2.173233e-02 2.096710e-02 1.577545e-02 2.208164e-02

                              ~~~~~~~~~~~~~~~~~~~~~~~~~~~~~~~~~~~
```

measure: reconfiguration speed 2^nd^ eigenvector

```
Nonparametric ANOVA (kruskal wallis) p-value: 2.089630e-01

      Source             SS            df             MS             Chi-sq       Prob>Chi-sq
    ___________    ______________    _______    ______________    ____________    ____________

    {'Columns'}    {[5.4905e+03]}    {[  3]}    {[1.8302e+03]}    {[  4.5376]}    {[  0.2090]}
    {'Error'  }    {[1.3850e+05]}    {[116]}    {[1.1940e+03]}    {0×0 double}    {0×0 double}
    {'Total'  }    {[    143990]}    {[119]}    {0×0 double  }    {0×0 double}    {0×0 double}


T1 vs T2  wilcoxon p value = 0.19152
        ---------
T2 vs T3  wilcoxon p value = 0.67328
        ---------
T3 vs T4  wilcoxon p value = 0.95899
        ---------

means:2.449856e-01 2.519729e-01 2.543254e-01 2.533717e-01
stds:1.776172e-02 1.282094e-02 1.531508e-02 1.304118e-02
```
